# A whole-brain monosynaptic input connectome to neuron classes in mouse visual cortex

**DOI:** 10.1101/2021.09.29.459010

**Authors:** Shenqin Yao, Quanxin Wang, Karla E. Hirokawa, Benjamin Ouellette, Ruweida Ahmed, Jasmin Bomben, Krissy Brouner, Linzy Casal, Shiella Caldejon, Andy Cho, Nadezhda I. Dotson, Tanya L. Daigle, Tom Egdorf, Rachel Enstrom, Amanda Gary, Emily Gelfand, Melissa Gorham, Fiona Griffin, Hong Gu, Nicole Hancock, Robert Howard, Leonard Kuan, Sophie Lambert, Eric Kenji Lee, Jennifer Luviano, Kyla Mace, Michelle Maxwell, Marty T. Mortrud, Maitham Naeemi, Chelsea Nayan, Nhan-Kiet Ngo, Thuyanh Nguyen, Kat North, Shea Ransford, Augustin Ruiz, Sam Seid, Jackie Swapp, Michael J Taormina, Wayne Wakeman, Thomas Zhou, Philip R. Nicovich, Ali Williford, Lydia Potekhina, Medea McGraw, Lydia Ng, Peter A. Groblewski, Bosiljka Tasic, Stefan Mihalas, Julie A Harris, Ali Cetin, Hongkui Zeng

## Abstract

Identification of the structural connections between neurons is a prerequisite to understanding brain function. We developed a pipeline to systematically map brain-wide monosynaptic inputs to specific neuronal populations using Cre-driver mouse lines and the recombinant rabies tracing system. We first improved the rabies virus tracing strategy to accurately identify starter cells and to efficiently quantify presynaptic inputs. We then mapped brain-wide presynaptic inputs to different excitatory and inhibitory neuron subclasses in the primary visual cortex and seven higher visual areas. Our results reveal quantitative target-, layer- and cell-class-specific differences in the retrograde connectomes, despite similar global input patterns to different neuronal populations in the same anatomical area. The retrograde connectivity we define is consistent with the presence of the ventral and dorsal visual information processing streams and reveals further subnetworks within the dorsal stream. The hierarchical organization of the entire visual cortex can be derived from intracortical feedforward and feedback pathways mediated by upper- and lower-layer input neurons, respectively. This study expands our knowledge of the brain-wide inputs regulating visual areas and demonstrates that our improved rabies virus tracing strategy can be used to scale up the effort in dissecting connectivity of genetically defined cell populations in the whole mouse brain.

## Introduction

The identity and function of neurons are determined not only by the inherent molecular and physiological characteristics of individual cells, but also by the synaptic connectivity through which diverse neuronal types form circuits. Advances in electron microscopy (EM) have enabled the reconstruction of synaptic resolution connectomes with different complexity, from the brain of *C. elegans* with 302 neurons to that of adult *Drosophila melanogaster* and larval zebrafish with ∼100,000 neurons^1–7^. Although brain-wide connectomics at single-cell resolution is currently beyond our grasp for complex nervous systems with over millions of neurons, different strategies have been applied to reveal the connectivity at distinct levels of resolution. Large-scale EM has been applied to reconstruct sub-volumes of the mouse and human brains, revealing both cellular and sub-cellular structures^8–12^. Whole-brain imaging of genetically labeled neurons can reveal the morphology of entire neurons and the fine details of dendritic and axonal coverage^13, 14^. Electrophysiological strategies such as multiple-patch clamp recordings reveal neurons that are synaptically connected and functionally depend on each other^15, 16^. Optogenetic activation of axon terminals of presynaptic neurons coupled with whole-cell recording of postsynaptic neurons have been utilized to examine neural connectivity over a range of spatial scales^16–20^. Wide-field imaging of genetically encoded calcium indicators allows simultaneous activity-monitoring of hundreds of neurons, which do not have to be synaptically connected^21^. Nonetheless, due to current technical limitations, these strategies cannot be used to reveal whole-brain connectivity in complex nervous systems.

Systematic mapping of afferent connectivity to specific cell populations has been greatly aided by the introduction of the monosynaptic, retrograde trans-synaptic rabies virus system^22, 23^. Rabies glycoprotein (RG)-deleted rabies viruses can be coupled with various genetic and viral tools to ensure the cell-type specific labeling of direct presynaptic inputs^24–36^. Many efforts have been made to improve the efficiency and specificity of rabies virus tracer while reducing its toxicity, including the construction of recombinant rabies viruses from the CVS N2c virus strain^37^, utilization of an engineered RG^25^, and generation of a double-deletion-mutant rabies virus^38^ and a self-inactivating rabies virus^39^. In addition, an intersectional rabies tracing strategy targeting Flp- and Cre-double labeled neurons has been generated to conduct cell-type-specific circuit tracing at an even more precise level^40^.

The ever-expanding repertoire of genetic and viral tools has enabled the construction of brain-wide mesoscale connectomes in a reasonable time frame^41–48^. In our effort to build the Allen Mouse Brain Connectivity Atlas, we combined viral tools, transgenic mouse lines, high-throughput imaging, and informatics to map brain-wide efferent connections at the level of cell classes^49, 50^. By delivering recombinant adeno-associated viruses (AAV) with Cre-dependent expression of enhanced green fluorescent protein (EGFP) to target brain areas of Cre transgenic lines, we labeled axons from selective Cre^+^ neuronal classes and subclasses. Our informatics pipeline, which includes registration of image series to the Allen Mouse Brain Common Coordinate Framework (CCF) and automatic segmentation of fluorescent axonal projections^51, 52^, enabled the quantification and comparison of whole-brain projections across multiple regions and cell classes. The resulting high-resolution mesoscale projection maps provide the foundation for in-depth dissection of the logic of mouse brain connectivity.

Aiming to construct a complementary afferent map of mouse brain-wide connectivity, we now developed an improved version of the monosynaptic rabies virus tracing system and incorporated rabies-mediated presynaptic input mapping into our pipeline. Our system consists of a single AAV helper virus that allows the accurate identification of starter neurons and rabies viruses expressing nucleus-localized EGFP marker to facilitate automatic quantification of presynaptic inputs. In this study, we utilized the retrograde connectome pipeline to map a brain-wide, cell-class-specific, presynaptic connectome for the mouse visual cortex, including both primary and higher visual areas. Mouse visual cortex contains at least ten visuotopically organized cortical areas^53–56^. These visual areas are strongly interconnected to form a hierarchical network with two visual streams as revealed by anterograde tracing^49, 57^, similar to what have been known in the primates and cat^58^. In primates, visual cortical hierarchy were defined by feedforward and feedback connections via laminar distribution of the retrogradely labeled neurons^59^. It remains largely unknown whether visual hierarchy and streams in mouse can also be defined with retrograde tracing.

By applying the monosynaptic rabies tracing system to Cre driver mouse lines labeling different excitatory and inhibitory neuron subclasses^44, 49^, our results reveal quantitative target-, layer- and cell-class-specific differences in the retrograde connectomes, despite similar global input patterns. We find that the retrograde connectomes of the same cell classes in different target areas are more different from each other than the retrograde connectomes of different cell classes in the same target area. Layer (L)-specific features are also identified, for example, L4 neurons receive more thalamic inputs and fewer inputs from higher-order association cortical areas, whereas L6 neurons are the main targets of contralateral/callosal inputs. Our study confirms previous findings of the dorsal and ventral streams in the mouse visual cortex^57^ and further reveals distinct subnetworks in the medial and lateral parts of the dorsal stream. Finally, our previous study showed that the hierarchical organization among different areas of the mouse visual cortex can be derived from axon termination patterns in the anterograde connectomes^49^, here we demonstrate that it can also be derived from the retrograde connectomes independently, via the feedforward and feedback projections mediated by upper- and lower-layer input neurons, respectively.

## Results

### A pipeline for the mesoscale retrograde connectome

To systematically map the whole-brain presynaptic inputs to different cell classes, we established a standardized high-throughput pipeline based on our pipeline for projection mapping across the entire brain^49, 50^, including the following steps: virus production and specimen generation (**Figure 1a-b**), data acquisition and processing (**Figure 1c-e**), and post-informatics characterization (**Figure 1 f-g**). The monosynaptic cell-type-specific rabies tracing system consists of EnvA-pseudotyped and glycoprotein-deleted rabies virus (EnvA RV^dG^) expressing histone-tagged EGFP (H2B-EGFP) and AAV helper virus conditionally expressing dTomato, the EnvA receptor TVA and RG. This system was coupled with Cre-driver mouse lines to reveal inputs to defined cell classes or types. The AAV helper virus and the monosynaptic rabies tracer were sequentially injected to the same target site with a three-week interval, followed by imaging of the whole brain one week after rabies infection. In this study focused on the visual cortex, our target site identification was guided by intrinsic signal imaging (ISI) of the visual areas. Rabies-labeled brains were imaged using high-throughput serial 2-photon tomography (STPT) at every 100 μm, with a total of 140 images for each brain. Injection polygons were drawn based on the expression of dTomato from the AAV helper virus, and the centroids of the injection site polygons were later used to verify and assign the target site. Image series were processed in the informatics pipeline for automatic segmentation of signal and registration to the Allen Mouse Brain Common Coordinate Framework version 3 (CCFv3)^51^ for subsequent data analyses. Brain sections were collected after STPT and those around the injection sites were further immunostained to enhance the dTomato signal. Starter cells were quantified after confocal imaging of the stained sections. Rigorous manual quality control steps were conducted to exclude experiments with noticeable tissue damage, injection failure, imaging failure or segmentation errors.

**Figure 1.**
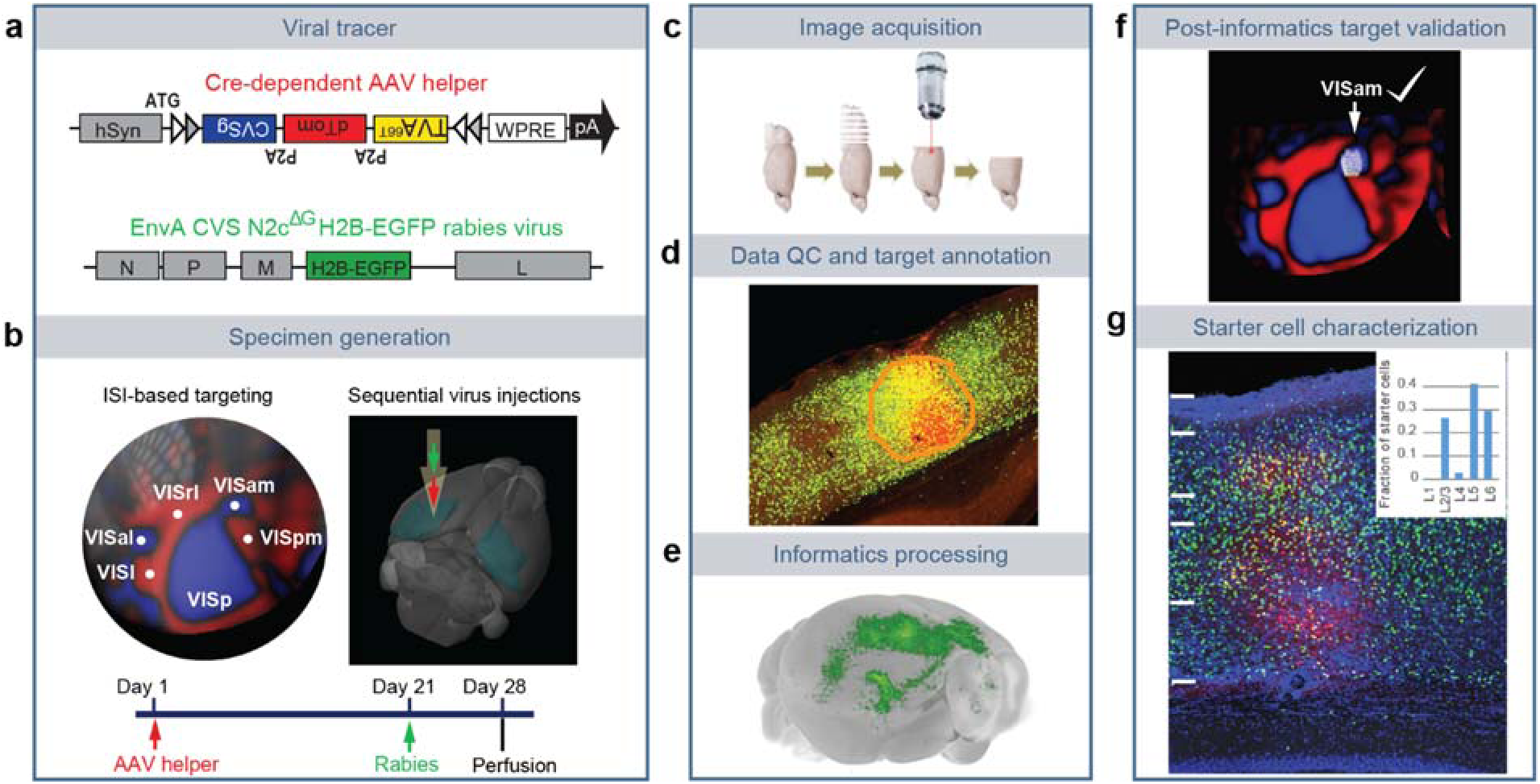
Pipeline identifying monosynaptic inputs to specific neuronal populations in the visual cortex. (**a**) Viral tools for mapping monosynaptic inputs to Cre-expressing neurons. The tricistronic AAV helper virus conditionally expresses TVA^66T^, dTomato, and rabies glycoprotein of the CVS N2c strain (CVSg) after Cre-mediated recombination. The EnvA-pseudotyped CVS N2c^dG^ rabies virus expresses histone-EGFP (H2B-EGFP) from the rabies G gene locus in the recombinant rabies virus genome. (**b**) ISI-based targeting and experimental timeline for virus injections and data analysis. (**c**) Sequential two-photon images were acquired at 100 μm interval and a total of 140 images were obtained for each brain. **(d)** Target sites were annotated by drawing injection polygons based on the expression of dTomato from the AAV helper virus. **(e)** Image series was automatically segmented and registered into the Allen CCFv3. **(f**) Injection site was verified post hoc by overlaying the injection site detected by AAV helper expression with an ISI image. **(g)** Starter cell characterization. Sections around the planned injection site were collected and dTomato signal was enhanced by immunostaining. Starter cells were then detected by co-expression of dTomato and nuclear EGFP, and the layer-distribution of the starter cells was analyzed.

We re-engineered several features of pre-existing rabies tracing tools to facilitate accurate identification of starter cells and automatic quantification of presynaptic inputs^31, 37, 60^ (**Figure 1a**). Our AAV helper virus uses the FLEX strategy to conditionally express a tricistronic cassette of TVA^66T^-P2A-dTomato-P2A-RG under the control of human synapsin promoter (hSyn), and the ATG of the tricistronic cassette was placed 5’ to the FLEX sites^60^ (**Figure 1a**). The co-expression of TVA, dTomato, and RG from the same expression cassette allows unambiguous identification of starter neurons. Both *in vivo* (**Supplementary Figure 1a-c** and **Supplementary Table 1**) and *in vitro* (not shown) tests of the new AAV helper virus demonstrate that this single AAV helper, with the use of the attenuated form of TVA, TVA^66T^, significantly reduces spurious virus labeling in the absence of Cre. Our rabies tracer is based on the CVS N2c^dG^ rabies strain^37^ and expresses H2B-EGFP. Compared with the EGFP-expressing and H2B-EGFP-experssing SAD B19^dG^ rabies viruses (**Supplementary Figure 1d-f** and **Supplementary Table 1**), the H2B-EGFP-expressing CVS N2c^dG^ viruses mediates stringent nucleus labeling, which facilitates automatic quantification of presynaptic cells by minimizing neurite labeling. This is the virus that we use throughout the paper and will refer to it simply as RV-H2B-EGFP.

We performed several control experiments to verify the specificities of our rabies virus tracing system in the absence of Cre, in the absence of RG provided by AAV helper viruses or in transgenic lines in which Cre is expressed in non-neuronal cell types. We found that applying the monosynaptic rabies tracing system to wild-type mice (*i.e.*, in the absence of Cre) led to only a few H2B-EGFP-labeled cells in the injection site, but no starter cells in the injection site and no H2B-GFP-labeled cells outside the injection site (**Supplementary Figure 2a-b,** and **Supplementary Table 1**). This shows that our system does not have the issue of spurious local rabies virus uptake due to low-level expression from the AAV helper in the absence of Cre^29, 32, 34, 61^, or local infection by small quantities of RG-coated RV^dG^ virus particles that may be present in the EnvA-pseudotyped rabies virus preparation.

We then confirmed that the trans-synaptic transfer of the recombinant rabies relies on the expression of rabies G from the AAV helper. A G-minus version of the AAV helper virus, which conditionally expresses TVA^66T^ and dTomato after Cre-mediated recombination, was injected into Cre^+^ mice, followed by the injection of rabies virus three weeks later. We observed H2B-EGFP-labeled cells only at the injection site and nowhere else in the brain^27^. This finding confirms that the presynaptic labeling is specific for the Cre^+^ starter cells expressing the tricistronic cassette and infected with RV-H2B-GFP rabies viruses.

Finally, we investigated whether rabies infection of non-neuronal cell types affects the specificity of monosynaptic tracing (**Supplementary Figure 2c-e** and **Supplementary Table 1**). We tested the monosynaptic rabies tracing system in three non-neuronal Cre lines, Olig2-Cre^62^, Tek-Cre^63^, and Aldh1l1-CreERT2^64^, which express Cre in oligodendrocytes, vascular endothelium, and astrocytes, respectively. Among all experiments using the non-neuronal Cre lines, with either the hSyn-driven AAV helper virus used in the pipeline or a similarly constructed CMV-driven helper virus, sporadic long-distance H2B-EGFP-labeled cells were found only in 50% of the injected Aldh1l1-CreERT2 mice (**Supplementary Figure 2e**). Our results show that the occasionally infected non-neuronal cells do not support the spread of rabies virus to neurons in local or distant areas.

Through rigorous testing, we show that our rabies tracing strategy presents minimal non-specific labeling in the absence of Cre and enables unambiguous identification of starter cells and automatic quantification of presynaptic inputs. By combining Cre-driver lines and the improved rabies tracing system, we aim to utilize our standardized pipeline for monosynaptic retrograde mapping to generate a comprehensive and quantitative mesoscale input network registered into a common 3D space.

### Comprehensive mapping of inputs to visual areas by cell types

We utilized our retrograde connectome pipeline to systematically map the presynaptic inputs of neurons in the visual areas by layers and cell classes. A total of 249 experiments across nine excitatory neuron Cre lines and five interneuron Cre lines were included in this study (**Figure 2a,** and **Supplementary Tables 2-3**). The Cre lines included those previously used in the Allen Mouse Connectivity Atlas to identify the organization of cortical connections in the mouse brain^49^. The nine excitatory neuron lines used were the pan-layer Emx1-IRES-Cre, L2/3 IT (Sepw1-Cre_NP39), L2/3/4 IT (Cux2-IRES-Cre), L4 IT (Nr5a1-Cre), L5 IT (Tlx3-Cre_PL56), L5 ET (A93-Tg1-Cre), L5 IT ET (Rbp4-Cre_KL100), L6 CT (Ntsr1-Cre_GN220), and L6b (Ctgf-2A-dgCre). The five interneuron lines used were Gad2-IRES-Cre, Ndnf-IRES2-dgCre, Pvalb-IRES-Cre, Sst-IRES-Cre, and Vip-IRES-Cre. All injections were performed into the left hemisphere, and injection site was verified post-hoc (**Methods**). We found that all but one experiment (which targeted the temporal association area, TEa), successfully targeted the visual areas (jointly labeled as VIS), including primary visual cortex (VISp), and higher visual areas (HVAs) such as lateral visual area (VISl, “LM”), posteromedial visual area (VISpm, “PM”), anteromedial visual area (VISam, “AM”), anterior area (VISa, “A”), anterolateral area (VISal, “AL”), rostrolateral visual area (VISrl, “RL”), and laterointermediate area (VISli, “Li”). Fourteen experiments targeted a subarea of the primary somatosensory area barrel field (SSp-bfd) bordering VISrl, which corresponds to previously defined VISrll region and displays extension of retinotopic organization lateral to VISrl^54^, and thus we refer to this barrel field subarea as SSp-bfd-rll. Locations of all injection centroids were plotted onto the top-down-view of the CCFv3 cortical map (**Figure 2b**).

**Figure 2.**
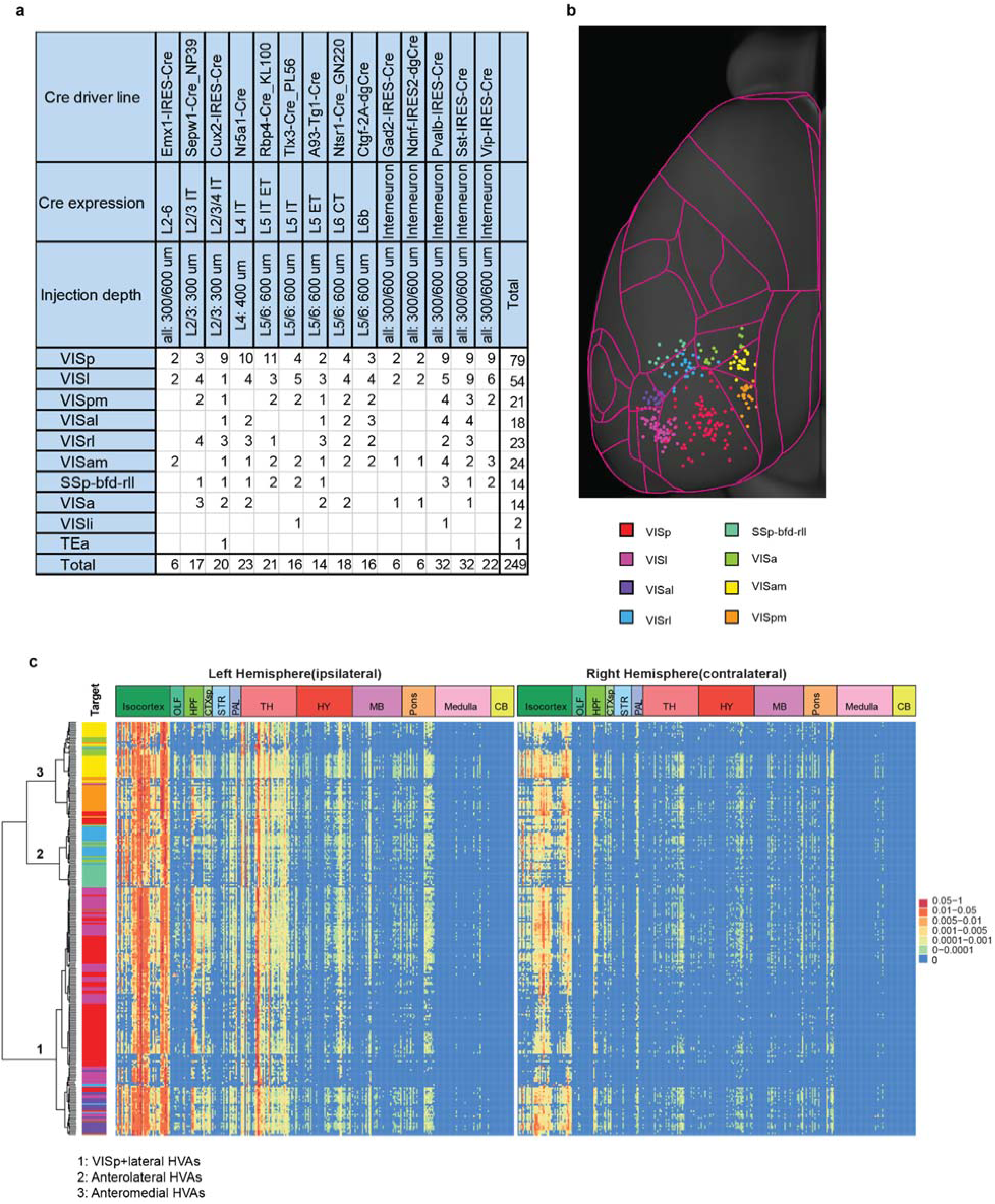
Identification of monosynaptic inputs to Cre-labeled neuronal classes in different visual areas. **(a)** Summary of Cre mouse lines, target areas, and numbers of the 249 experiments in the visual areas. The injection target areas were verified based on overlay of injection site polygons with ISI images or the position of injection site polygons in the Allen CCFv3. **(b)** Mapping of each injection centroid in the top-down view of mouse cortex. Color indicates different visual areas. Red: VISp; Magenta: VISl; Purple, VISal; Light blue: VISrl; Aquamarine: SSp-bfd-RLL; Light green: VISa, Yellow: VISam; Orange: VISpm. **(c)** Matrix showing normalized inputs from the ipsilateral (left) hemisphere and the contralateral (right) hemisphere for the 249 experiments. Each row represents one of the 249 experiments; columns are ordered by 12 major brain divisions; rows are organized according to hierarchical clustering of the input patterns. The input signal per structure was measured by the informatics data pipeline and represented by per structure input signal volume (sum of detected signal in mm^3^). To reduce false positive signal, we identified a set of 92 negative brains that were processed through the pipeline, but showed no rabies-mediated GFP expression, and used this negative dataset to calculate the threshold of false positive signal, i.e., the value of mean input signal volume plus 6 standard deviations for each of the 314 ipsilateral and 314 contralateral major structures of the brain. Input signal not passing this threshold was set to “0”. A manually validated binary mask was then applied to remove artifacts in informatically-derived measurements of input signal. Following these two steps, input signal volume of a given structure was normalized to the total input of the brain. Value in each cell of the matrix represents the input signal volume in the given brain area as the fraction of total input of the brain. Color in the “Target” represents the verified injection target area of the experiment in each row, as color-coded in (b).

We quantified presynaptic inputs in each brain area using an automated image segmentation algorithm trained to detect the fluorescence signal from nucleus-localized H2B-EGFP^52^. We validated the accuracy of our automatic signal detection and quantification of inputs following registration to the CCFv3 by comparing the informatically measured per structure input signal volume (sum of detected signal in mm^3^) with manual counting of labeled cells (**Supplementary Figure 3a-g**). In the six structures from the cortex, thalamus, and cortical subplate, strong positive linear correlations were found between automatic measurement and manual counts (R^2^ in the 0.62-0.98 range). Therefore, for subsequent analysis, we used the automatically calculated input signal volume for each structure.

We find that expression of dTomato from the AAV helper virus faithfully reflects the presence of Cre recombinase. The starter cells show distinct layer-specific distribution patterns consistent with the Cre expression of the respective transgenic lines. The numbers of starter neurons vary between Cre lines and between different experiments within the same Cre line (**Supplementary Figure 3h-i,** and **Supplementary Table 2**). Although the overall presynaptic labeling signal increases with the number of starter cells, there is not a strong linear correlation (R^2^ = 0.54) between the number of starter cells and total input signal volume within the brain (**Supplementary Figure 3j**). It suggests that postsynaptic cells may receive convergent input from presynaptic cells, which in turn can make divergent connections to different postsynaptic cells. Previously, we compared the whole-brain projections across animals by normalizing the projection signals to the size of the infection area^49^. Here, due to the lack of strong linear correlation between whole brain input signal and the number of starter cells in the injection site, we instead use the fraction of whole brain inputs as our measure of connectivity strength per region, i.e., the input signal volume per brain structure divided by the input signal volume of the entire brain.

We next constructed a brain-wide matrix for inputs to the visual areas, focusing on the fraction of whole brain inputs from 314 major structures at a mid-ontology level from the CCFv3 (**Figure 2c,** and **Supplementary Table 3**). At this anatomical level, hierarchical clustering of the fraction of whole brain inputs measured from the 249 experiments separate all experiments into three major clusters that are correlated with the spatial proximity of the injection sites: the VISp and lateral HVA cluster (VISl and VISal), the anterolateral cluster (SSp-bfd-rll, and VISrl), and the anteromedial cluster (VISam, VISpm, and VISa). We find that compared to the brain-wide output projections from mouse visual areas^49^, presynaptic inputs come from a broader collection of brain areas (**Figure 2c**). Visual areas receive the strongest inputs from isocortex (fraction of inputs: median: 0.82, range: 0.52-0.91), followed by thalamus (median: 0.13, range: 0.06-0.46) and hippocampal formation (HPF) (median: 0.02, range: 0-0.16) (**Supplementary Figure 3k)**.

### Brain-wide inputs to the VISp and HVAs

Given the correlation between whole brain input patterns and spatial proximity of the injection sites, we compared the presynaptic inputs across all experiments in VISp,VISl, VISal, VISrl, VISa, VISpm, VISam and SSp-bfd-rll. This was possible due to the use of precise injections guided by ISI and sufficient coverage and numbers of injections for all visual areas. Comparison of bilateral inputs from the whole brain (isocortical modules^49^, olfactory area (OLF), HPF, cortical subplate (CTXsp), striatum (STR), pallidum (PAL), thalamus (TH), hypothalamus (HY), midbrain (MB), pons, medulla (MY), and cerebellum (CB)) to the eight target areas reveal overall similar, but quantitatively different global input patterns to different targets, with dominant inputs from the isocortical modules and thalamus (**Figure 3a**). Visual areas also receive strong presynaptic inputs from HPF, which are mainly found in lateral entorhinal cortex (ENTl), medial entorhinal cortex (ENTm), CA1 and the post-, pre- and parasubiculum (POST, PRE, PAR) (**Supplementary Figure 4**). In primate, the hippocampal complex and entorhinal cortex are placed at the top of the visual area hierarchy^58^. The entorhinal cortex serves as the interface for a multi-synaptic pathway connecting the visual area with the hippocampus, in which ENTl conveys ventral-stream input to the hippocampus and ENTm conveys dorsal-pathway input^65, 66^. Our results reveal direct entorhinal and CA1 inputs to the visual cortex, indicating a visual cortical-hippocampal-visual cortical loop of information processing. A bias for ventral CA1 input to the visual area is also observed, consistent with the distinct projection pathways from the ventral and dorsal CA1^67^.

**Figure 3.**
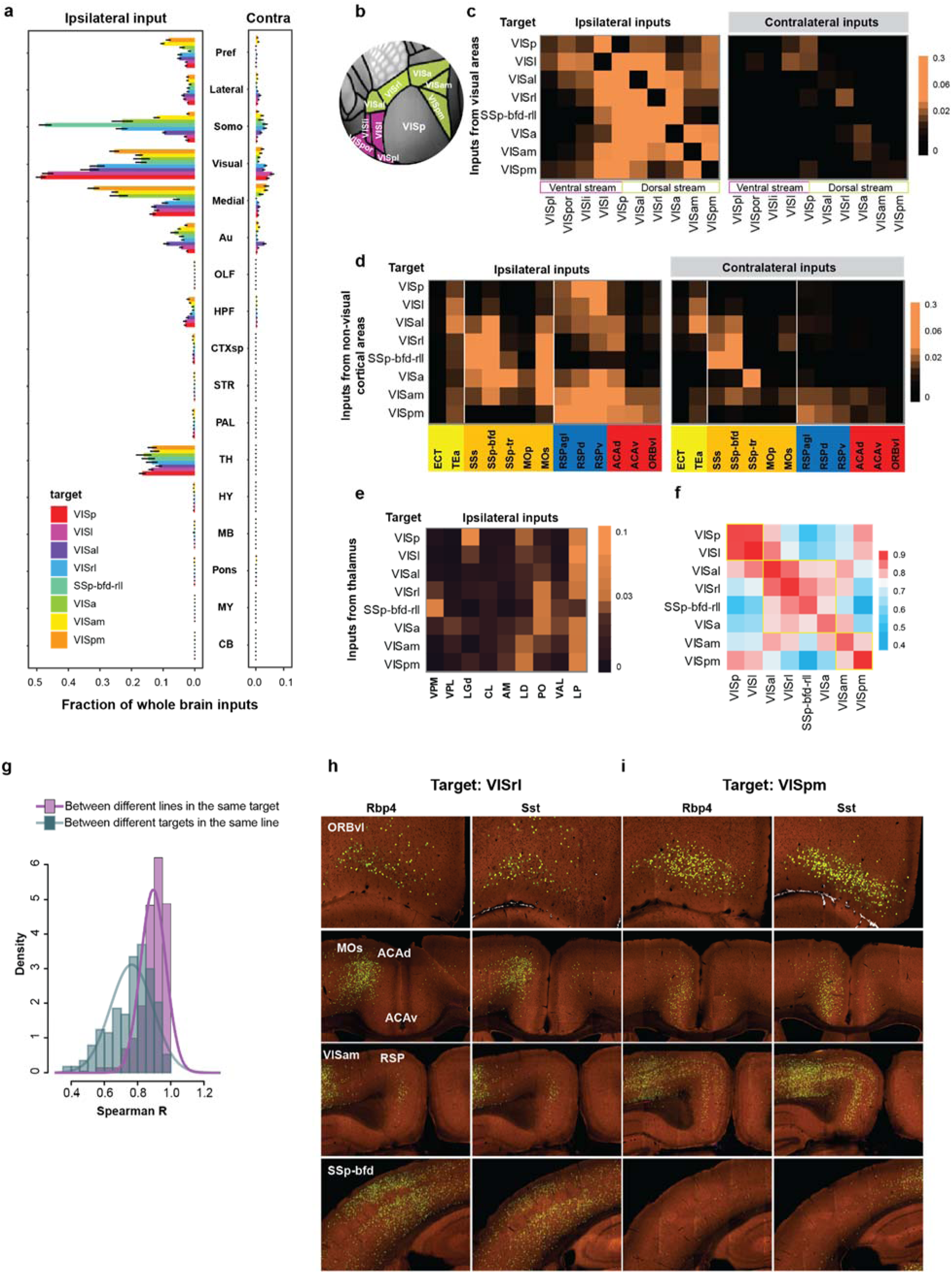
Comparison of whole-brain inputs to neurons in the primary visual cortex and higher visual areas. **(a)** Comparison of inputs from the isocortex, olfactory area (OLF), hippocampal formation (HPF), cortical subplate (CTXsp), striatum (STR), pallidum (PAL), thalamus (TH), hypothalamus (HY), midbrain (MB), pons, medulla (MY), and cerebellum (CB) to the visual areas. Inputs from the isocortex were divided into six modules: prefrontal (Pref), lateral, somatomotor (Somo), visual, medial, and auditory (Au) (data shown as mean ± s.e.m.). **(b)** Visual areas in the ventral and dorsal streams in the top-down view of the CCFv3 cortical map. **(c-e)** Matrices showing the inputs to visual area targets received from within the visual cortex **(c)**, non-visual isocortical modules **(d)**, and thalamus **(e)**. Each row represents the average per structure normalized inputs for experiments with starter cells confined to the same target area, and each cell within the matrix represents the mean value of normalized inputs from a given source area. Visual areas are separated into the dorsal and ventral streams and higher visual areas are in a clockwise order from VISp. Cortical areas with noticeable inputs to visual areas are selected and ordered by module membership. Thalamic areas with noticeable inputs to visual areas are selected and ordered by hierarchical orders. Areas in the thalamus, sensory-motor cortex related, are highlighted in pink and areas in the thalamus, polymodal association cortex related, are highlighted in blue. **(f)** Matrix showing Spearman’s correlation coefficients (R) between experiments within the same target (intra-target) and between experiments across different targets (inter-target) using the input dataset in c-e. Areas with high correlations are highlighted in yellow boxes. The intra-target correlation was calculated as the mean of the spearman correlation coefficients between experiments from different Cre-defined cell types, while the inter-target correlation was calculated as the mean of the spearman correlation coefficients between experiments of the same Cre-defined cell types in different targets. **(g)** Density plot of Spearman’s R for intra-target experiments of different cell types and inter-target experiments of same cell types using the input dataset in b-d. The mean of the intra-target Rs is greater than the inter-target Rs, suggesting that experiments of different cell types in the same target are highly similar to each other. **(h-i)** Comparison of retrograde input patterns between injections in VISrl and VISpm of Rbp4-Cre and Sst-Cre lines. Rbp4 and Sst experiments in VISrl both show strong inputs from ACAd and SSp-bfd, and weak inputs from RSP and VISam. In contrast, Rbp4 and Sst experiments in VISpm show strong inputs from ACAv, RSP, ORBvl, and VISam, and weak inputs from ACAd and SSp-bfd.

Inputs from other anatomical structures each provide less than 1% of whole brain inputs, and the fraction of inputs span more than three orders of magnitude, ranging from claustrum (CLA) in CTXsp, diagonal band nucleus (NDB) in PAL, caudoputamen (CP) in STR and lateral hypothalamic area (LHA) in HY each representing ∼0.1% of whole brain inputs, globus pallidus, external segment (GPe) in PAL, basolateral amygdalar nucleus (BLA) in CTXsp and zona incerta (ZI) in HY each accounting for ∼0.01% of whole brain inputs, dorsal peduncular area (DP) in OLF, locus ceruleus (LC) in pons, and superior colliculus (SC) in MB each accounting for ∼0.001% of whole brain inputs, to areas in MY each accounting for ∼0.0001% of whole brain inputs (**Supplementary Figure 4**). CLA is reciprocally connected to various sensory-related brain areas^68^, and the observed strong CLA input to the visual cortex suggests a possible role of CLA in integrating visual processing with other sensory cues. Sparse SC inputs are found in only 11% of all experiments (**Supplementary Table 3**) in accordance with the major relay of SC visual input via the lateral posterior nucleus (LP) of the thalamus. Rare inputs in several structures of MY are also found in less than 10% of all experiments (**Supplementary Table 3**), which could be missed using other connectivity mapping techniques.

The distribution of subcortical inputs strongly suggests the involvement of neuromodulatory systems in regulating the mouse visual cortex. Clustered inputs are found in NDB (fraction of whole brain inputs in NDB > 0 in 93% of all experiments), and substantia innominata (SI) (fraction of whole brain inputs in SI > 0 in 75% of all experiments) (**Supplementary Figure 4 and Supplementary Table 3),** consistent with the innervation of the visual cortex by forebrain cholinergic neurons^69, 70^. LC primarily consists of noradrenergic neurons, which send outputs to broad regions of the brain. Our results reveal sparse LC inputs to both VISp and HVAs (fraction of whole brain inputs in LC > 0 in 66% of all experiments) (**Supplementary Figure 4 and Supplementary Table 3)**, suggesting that the visual cortex is part of the ascending LC efferent pathway innervating the limbic system, midbrain, thalamus, basal forebrain and neocortex^71, 72^. In the dorsal raphe (DR) where dorsal cortex-projecting serotonin neurons were previously identified^73^, sparsely labeled presynaptic neurons (∼0.01% of whole brain inputs) are found in 63% of all experiments (**Supplementary Figure 4 and Supplementary Table 3)**. Identification of potential monosynaptic projections from neuromodulator-expressing neurons to both VISp and HVAs suggests that locally released neuromodulators can affect all levels of visual processing.

We then focused on individual cortical and thalamic source areas and used the average from all experiments (including all Cre lines) in the same target to compare the input strength from a given presynaptic brain structure to the eight visual targets (**Figure 3b-e**). The input patterns strongly suggest the presence of two subnetworks equivalent to the dorsal and ventral cortical streams in primates^74^. The nodes in the dorsal stream and the ventral stream of the mouse visual system were previously mapped based on the anterograde projection strength, with VISl, VISli, VISpl, and VISpor in the ventral stream and VISal, VISrl, VISa, VISpm and VISam in the dorsal stream^57^. Our study covers VISal, VISrl, VISa, VISpm and VISam in the dorsal stream, which tends to receive more inputs from each other and fewer inputs from ventral stream structures such as VISpl, VISpor, and VISli (**Figure 3c**). Consistent with a strong correlation between input patterns and starter cell locations, we find that two adjacent areas, SSp-bfd-rll and VISrl, show similar input patterns characteristic of dorsal stream structures. Both receive strong inputs from the somatomotor cortex, including the secondary somatosensory cortex (SSs), primary somatosensory area barrel field (SSp-bfd), and secondary motor (MOs), but SSp-bfd-rll receives fewer inputs from the midline cortical areas such as anterior cingulate area (ACA) and retrosplenial (RSP) as compared to VISrl (**Figure 3d**). Given the input patterns of SSp-bfd-rll, we place SSp-bfd-rll as part of the dorsal stream pathway. The ventral stream node, VISl, in general presents similar input pattern as VISp. However, VISp can be distinguished from VISl and other HVAs based on the preference for inputs from the dorsal lateral geniculate complex (LGd) of the thalamus over LP (**Figure 3e**).

Within the dorsal stream, we also find that anterolateral structures such as VISal, VISrl, and SSp-bfd-rll present strong inputs among themselves, while receiving relatively few inputs from the medial structures such as VISam and VISpm (**Figure 3c**). In contrast, VISam and VISpm show strong mutual connections, while VISpm receives relatively few inputs from the anterolateral structures (**Figure 3c**). In addition, comparison between inputs to the anterolateral and medial structures reveals that the anterolateral structures of the dorsal stream receive stronger inputs from the somatomotor module, whereas the medial structures, VISam and VISpm receive stronger inputs from RSP, ACA and ventrolateral orbital area (ORBvl) (**Figure 3d**). The anterolateral and medial structures of the dorsal stream also present distinct thalamic input patterns, with VISam and VISpm receiving strong inputs from the anteromedial nucleus (AM) and lateral dorsal nucleus (LD) and anterolateral structures receiving strong inputs from the posterior complex (PO) and ventral anterior-lateral complex (VAL) (**Figure 3e**). Such input patterns can be observed in both excitatory neuron and interneuron experiments (**Supplementary Figure 5**). To evaluate the similarity of input patterns between different targets, we calculated Spearman’s correlation coefficients (R) for experiments within the same target and for experiments across different targets (**Figure 3f**). Our results suggest the possibility of two subnetworks in the dorsal stream, one consisting of medial structures of VISam and VISpm and the other consisting of anterolateral structures of VISal, VISrl, SSp-bfd-rll and VISa, with a gradual transition between the two subnetworks from medial to lateral.

Given the unique input pattern to each visual area, we further investigated the effects of different starter cell classes on the input patterns. We find that the correlation between input patterns of different cell classes in the same target is higher than that of the same cell class in different targets (**Figure 3g-i**). Using experiments in VISrl and VISpm as the example, regardless of the starter cell classes, VISrl receives characteristically stronger inputs from ACAd, MOs, and SSp-bfd than VISpm, whereas VISpm receives stronger inputs from ORBvl, ACAv, and RSP than VISrl. Our results suggest that the presynaptic input patterns, as quantified by our tracing system, are predominantly determined by the spatial location of the starter cells, and that different cell classes in the same target receive similar global input patterns.

### Comparison of brain-wide inputs to excitatory neurons in different layers

We next focused on quantifying the brain-wide inputs to different Cre-defined glutamatergic excitatory neuron subclasses within a single target area, VISp. Despite the variation in starter cell numbers and layer distributions (**Supplementary Figure 6a,b,d**), the overall input patterns for any specific location in VISp are similar between different Cre driver lines (**Supplementary Figure 6c,e**), with most inputs arising from isocortex, followed by thalamus and HPF. Compared to other layer-specific lines, the L4 line receives significantly more input from the thalamus (*P* < 0.001, Tukey multiple comparisons of means), consistent with the notion that feedforward signal from the visual thalamus is mostly received by L4 neurons in the visual cortex^75–77^.

We constructed an input strength matrix for experiments targeting 9 Cre-defined excitatory neuron populations in VISp (**Figure 4a and Supplementary Figure 7a**), and compared that to the brain-wide output projection matrix (**Supplementary Table 4**) for the same Cre-defined neuron populations in VISp (**Figure 4b and Supplementary Figure 7b**). In each hemisphere, 43 cortical structures are organized into six corticocortical connectivity modules: prefrontal, lateral, somatomotor, visual, medial, and auditory, as revealed by projection connectivity in our previous study^49^. We find that VISp receives the strongest input from areas within the visual module, followed by visual areas within the medial module. Outside these areas, VISp excitatory neurons receive the majority of inputs from ACA and ORB in the prefrontal module, TEa and ectorhinal (ECT) areas in the lateral module, SSs, SSp-bfd, and MOs in the somatomotor module, RSP in the medial module, and the auditory module. Within the prefrontal module, significantly more presynaptic neurons to VISp are found in ACAd (dorsal part) than ACAv (ventral part, paired t-test, *P* < 0.001) and in ORBvl (ventrolateral part) than ORBm (medial part) and ORBl (lateral part, paired t-test, *P* < 0.001). The striking similarity between the patterns of VISp intracortical input and output reveals the reciprocity of corticocortical connections (**Figure 4b**).

**Figure 4.**
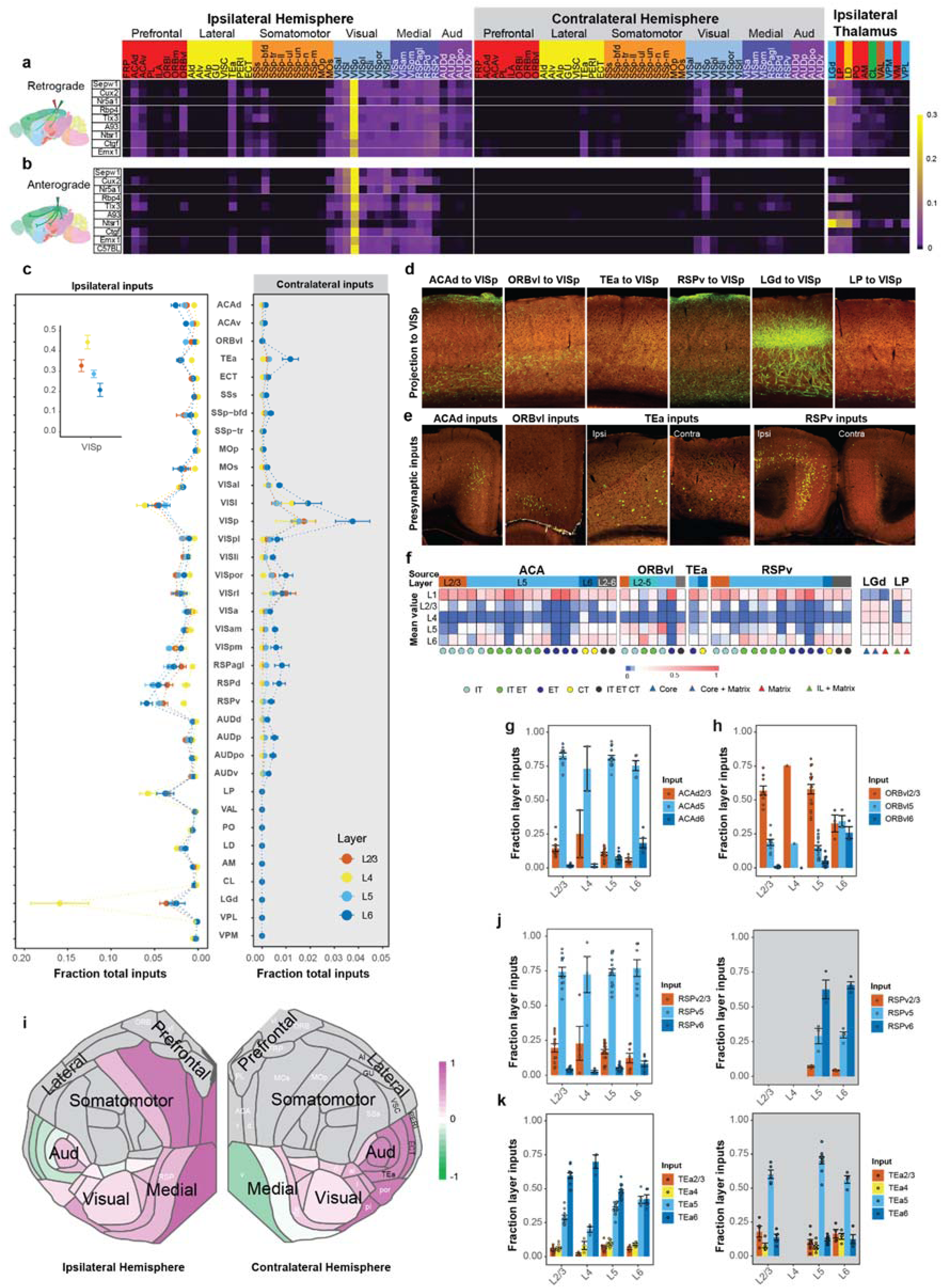
Comparison of the whole-brain input patterns to excitatory neuron subclasses in different layers of the primary visual cortex. **(a)** Matrix showing normalized inputs from the ipsilateral and contralateral isocortex, and ipsilateral thalamus to excitatory neurons in different layers of VISp. Each row of the matrix represents the mean normalized per structure input signals for experiments in each Cre line. Rows are organized based on layer-specific distribution of the starter cells. The 43 ipsilateral (left) and 43 contralateral (right) cortical areas are ordered first by module membership (color coded) then by ontology order in the Allen CCFv3. The ten thalamic nuclei are ordered based on the strength of inputs and are colored by the thalamocortical projection classes (blue: core, green: intralaminar, brown: matrix-focal, and red: matrix-multiareal). **(b)** Matrix showing normalized axonal projections from VISp to the ipsilateral and contralateral isocortex, and ipsilateral thalamus shown in (a). Anterograde tracing experiments (**Supplementary Table 4**) from the Cre mouse lines used in (a) and C57BL/6J were included, and rows represent the mean normalized per structure projection signals for experiments in each mouse line. **(c)** Comparison of ipsilateral and contralateral inputs from cortical areas and thalamic nuclei to excitatory neurons in different layers of VISp. Data are shown as mean ± s.e.m. **(d)** Representative STPT images showing laminar termination patterns of axon projections in VISp from higher-order association cortical areas and thalamic nuclei. **(e)** Laminar distribution patterns of presynaptic input cells in higher-order association cortical areas that project to VISp. **(f)** Normalized laminar termination patterns in VISp for projections from higher-order association cortical areas and thalamic nuclei. Each column represents the relative projection strengths by layer for a unique combination of Cre-defined cell classes and source areas. The average value was taken when n > 1. L6b was excluded due to low accuracy in informatic quantification of projection signal in L6b. **(g-h)** Laminar distribution of long-range inputs from ACAd (g) and ORBvl (h) to excitatory neurons in different layers of VISp. The fraction layer input is calculated as the fraction of the total input signal in a given source area across layers. L1 is excluded from the analysis due to overall lack of signal in this layer. The calculated fraction layer inputs are consistent with representative images of inputs in ACAd and ORBvl to VISp (e). Data are shown as mean ± s.e.m. **(i)** Comparison of L5 and L6 preference for source cortical areas in the ipsilateral (left) and contralateral (right) hemispheres sending presynaptic inputs to VISp. The preference score for a given cortical area is calculated as (L5 input - L6 input) / (L5 input + L6 input). Each source cortical area was colored according to its preference score. **(j-k)** Laminar distribution of inputs in RSPv (j) and TEa (k) to VISp as examples of source cortical areas located in the medial (RSPv) or lateral (TEa) areas of the cortex. Data are shown as mean ± s.e.m.

Ten thalamic nuclei with the strongest inputs to VISp are included in the input strength matrix (**Figure 4a**). Three thalamic nuclei, LGd, LP and LD, collectively account for more than 70% of the thalamic inputs. These three nuclei also receive strong VISp projections from the L6 CT neurons labeled by the Ntsr1 Cre line as well as L5 ET neurons labeled by Rbp4 and A93 Cre lines (**Figure 4b**), with L6 CT mainly targeting LGd and L5 ET preferentially targeting LP and LD. The brain-wide monosynaptic input matrix also reveals that VISp receives strong inputs from ENTl, ENTm, PAR, POST, CLA and NDB (**Supplementary Figure 7a**).

We then compared the input patterns of excitatory neurons in different layers of VISp (**Figure 4a, c and Supplementary Figure 7a, c**). We find that L4 neurons receive the least amount of input from higher-order association cortical areas, including ACA, ORBvl, TEa and RSP, as well as from subcortical input areas, and receive significantly more inputs from LGd. In contrast, L2/3, L5 and L6 neurons generally receive more inputs from higher-order association cortical areas. Specifically, L5 neurons in VISp receive the most input from ORBvl, while L6 neurons in VISp receive the most input from ACAd and ACAv. This layer-specific input pattern is supported by the axonal lamination patterns from cortical and thalamic areas to VISp (**Figure 4d,f**), where ORBvl axons primarily ramify in L1 and L5 of VISp, ACAd axons mainly reside in L1 and L6, and axons from LGd mainly ramify in L4.

Subsequently, we analyzed the laminar distribution of cortical inputs to excitatory neurons in different layers (**Figure 4e, g-h**). We find that the presynaptic input neurons in higher-order association cortical areas are often located in deep layers (L5 and L6), regardless of the layer location of starter cells in VISp. For example, in ipsilateral ACAd, L5 contains the most presynaptic neurons compared to other layers, with a similar preference for L5 observed for inputs to excitatory neurons located in different layers of VISp (**Figure 4e,g**). A noticeable exception is ipsilateral ORBvl, in which VISp L2/3, L4 and L5 neurons preferentially receive inputs from L2/3 ORBvl whereas no preference in layer distribution is found for input to L6 neurons of VISp (**Figure 4e,h**).

We also investigated whether inputs from homotypic ipsilateral and contralateral cortical areas to VISp arise from different layers (**Figure 4i-k**) and which source layer contributes most to VISp. Since most inputs arose from L5 and L6, we calculated a preference score for a given cortical area as (L5 input - L6 input)/(L5 input + L6 input). We find that medial areas of the ipsilateral hemisphere show a bias toward L5 input to VISp, with the preference gradually shifting towards L6 in lateral areas, whereas areas of the contralateral hemisphere present an opposite bias: medial areas show L6 bias, and lateral areas show L5 bias (**Figure 4i**).

These distinct features in whole-brain input patterns to excitatory neurons in different layers of VISp can also be found in VISl (**Supplementary Figure 8 and Supplementary Table 5)**. Similar to our observation in VISp, we find that L4 overall receives more inputs from thalamic areas and fewer inputs from higher-order cortical areas, most inputs from higher-order cortical areas are from the deep layers, and ipsilateral and contralateral cortical areas present different laminar distribution of input neurons to the same target. We also find generally consistent input patterns to excitatory neurons in different layers of other HVAs as of VISp and VISl, though due to smaller number of experiments in each layer of each region (**Figure 2a**) we do not provide quantitative analysis here.

### Distinct presynaptic inputs to L6 excitatory neurons of visual areas

We observed that L6 CT and L6b neurons (as labeled by Ntsrt1 and Ctgf Cre lines) clearly receive more inputs from the contralateral cortex compared to excitatory neurons in the other layers (**Figure 4a and Supplementary Figure 6e**). These inputs mostly originate from the contralateral visual, medial, and auditory modules. To further investigate whether the layer distribution of starter neurons is the key factor in determining the level of contralateral inputs, we identified 89 experiments in VISp and HVAs with starter cells restricted to a single layer and compared the contralateral and ipsilateral inputs between the experiments. Overall, L6 neurons across VIS receive more contralateral inputs from all six isocortical modules than neurons in other layers (**Supplementary Figure 9**). Quantitative analysis suggests that the effect of layer on the ratio of contralateral to ipsilateral isocortical inputs is significant (two-way ANOVA, *P* < 0.001), and that the location of the target site (VISp or HVAs) does not significantly affect the ratio of contralateral to ipsilateral isocortical inputs (*P* = 0.37). Our results suggest that L6 of the visual area has distinct retrograde connectivity compared to the other layers.

### Brain-wide inputs to VISp interneurons

To explore presynaptic inputs to distinct GABAergic interneurons, we employed various Cre lines driven by genes corresponding to major interneuron subclasses: parvalbumin (Pvalb)-expressing, somatostatin (Sst)-expressing, vasoactive intestinal peptide (Vip)-expressing neurons, and neuron-derived neurotrophic factor (Ndnf)-expressing neurons which are mostly L1 neurogliaform cells. We also included the Gad2-Cre line to cover all cortical interneurons.

Despite variation in starter cell numbers (**Supplementary Figure 3**) and layer distribution, the overall global patterns are again similar between Cre lines, regardless of excitatory or inhibitory type (**Figure 5a**). We quantified the fraction of inputs from cortical modules and thalamus to excitatory neurons and interneuron cell classes located in VISl, VISp and VISam, where we had experiments covering almost all the 14 Cre-defined cell classes. We find that, compared to excitatory neurons, interneurons overall receive more inputs from thalamus and ipsilateral visual cortical module (**Figure 5b)**, and fewer inputs from contralateral cortical modules (**Figure 5b-c**), suggesting that intra-module inputs exert greater influence on interneurons than excitatory neurons.

**Figure 5.**
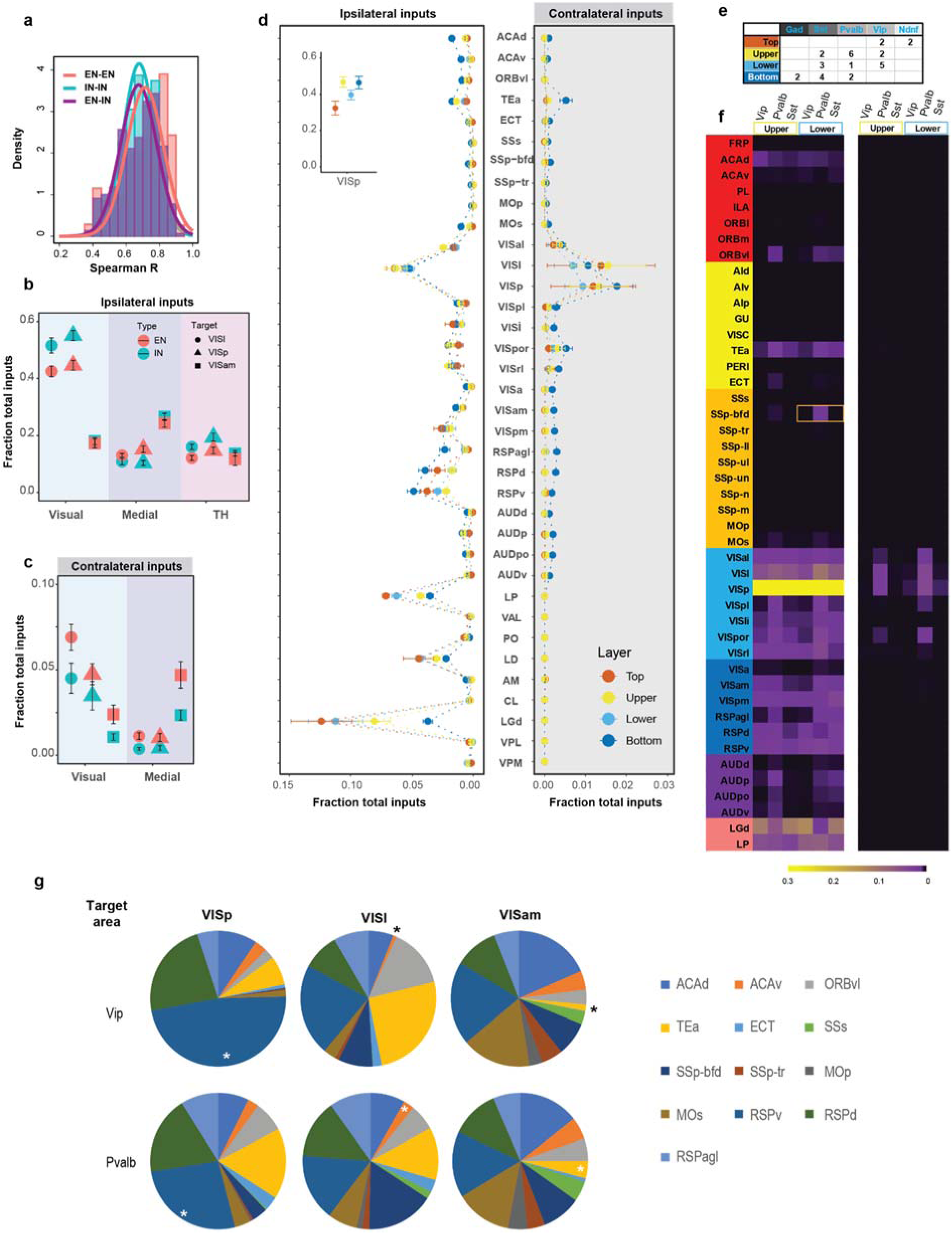
Comparison of the whole-brain input patterns to different interneuron subclasses in the primary visual cortex. **(a)** Density plot of Spearman’s R values for whole-brain input patterns to different excitatory neuron (EN) subclasses, those to different interneuron (IN) subclasses, and Rs measured between EN input patterns and IN input patterns. The means of the inter-cell-class Rs and intra-cell-class Rs are close to each other, suggesting that input patterns to EN and IN classes are highly similar to each other. **(b-c)** Comparison of ipsilateral (b) and contralateral (c) inputs from the visual and medial cortical modules and thalamus to EN and IN cell classes located in the VISl (circle), VISp (triangle), and VISam (square). Data are shown as mean ± s.e.m. **(d)** Comparison of inputs from ipsilateral and contralateral cortical areas and thalamic nuclei to interneurons located in different depths of VISp. Data are shown as mean ± s.e.m. **(e)** Summary of the depth distribution of IN experiments in VISp. **(f)** Matrices showing normalized inputs in the ipsilateral cortex, LP and LGd to the three IN subclass experiments in the upper and lower groups. Each column of the matrix represents the mean normalized input signals for the IN subclass. The area with statistically significant difference in inputs to Pvalb and Vip cells is boxed in yellow. **(g)** Comparison of relative input strength from higher-order cortical areas to Pvalb and Vip cells in VISp, VISl and VISam. Areas with statistically significant differences between the two interneuron subclasses are labeled with asterisks.

We then focused on different interneuron subclasses in VISp to investigate the possibility of cell-class-specific input patterns. To account for the contribution of layer distribution of starter cells to input patterns (**Supplementary Figure 10)**, we divided the 31 interneuron experiments in VISp into four different groups based on the depth of starter cell population: the Top group contained experiments with starter cells restricted to L2/3 (we found very few L1 starter cells), whereas the Upper, Lower, and Bottom groups had progressively more starter cells in deep layers (**Supplementary Figure 10b**). Although starter cells in these groups are rarely restricted to a single layer, we find distinct input patterns of interneurons, especially between the Bottom group and others. Compared to other groups, the Bottom group receives more inputs from higher-order cortical areas, including the frontal, sensorimotor, and auditory modules, and the fewest inputs from thalamic areas such as LGd and LP (**Figure 5d)**. Consistent with the observation of L6 excitatory neurons receiving extensive contralateral cortical inputs, the Bottom group also receives more contralateral cortical input than the other groups (**Figure 5d**). We then compared Sst, Vip, and Pvalb experiments in the Upper and Lower groups to avoid the confounding influence of layer distribution of starter cells on input patterns (**Figure 5e-f**). Despite variations of normalized inputs from higher-order cortical areas between Vip and Pvalb, statistical significance was not observed for most presynaptic areas, likely due to limited sample sizes. The distinct patterns of cortical inputs between Vip and Pvalb can also be observed when comparing all Vip and Pvalb experiments in VISp, VISl, and VISam, regardless of starter cell layer distribution **(Figure 5g)**. Both cell classes present unique target-specific cortical input patterns, and within the same target, Vip and Pvalb also differ in the relative input strength from selected higher-order cortical areas.

### Local inputs to excitatory neurons and interneurons

In experiments where the starter cells were restricted to a specific layer of VISp or VISl, we also examined local inputs (**Figure 6, Supplementary Figure 11**). We define the layer preference of any local input as the fraction of input in that layer compared to all local inputs. We find that the starter cells in each layer receive characteristic local input patterns. In VISp, L2/3 excitatory neurons preferentially receive inputs from L4 and L5. L4 excitatory neurons receive the fewest inputs from L2/3 and preferentially receive inputs from L4 and L5. L5 excitatory cells receive strong inputs from L2/3 to L6, with a noticeable preference for L2/3 input, and L6 excitatory neurons preferentially receive inputs from the deeper layers. With the exception of L2/3, starter cells receive dense inputs from other cells in the same layer. Our results support dense inputs from L4 to L2/3, despite weak input from L2/3 to L4, and dense reciprocal inputs between L2/3 and L5. Analysis of the local input patterns to excitatory neurons with layer-specific distribution in VISl also reveals that starter cells in each layer receive characteristic local inputs similar to excitatory neurons in VISp (**Supplementary Figure 11**).

**Figure 6.**
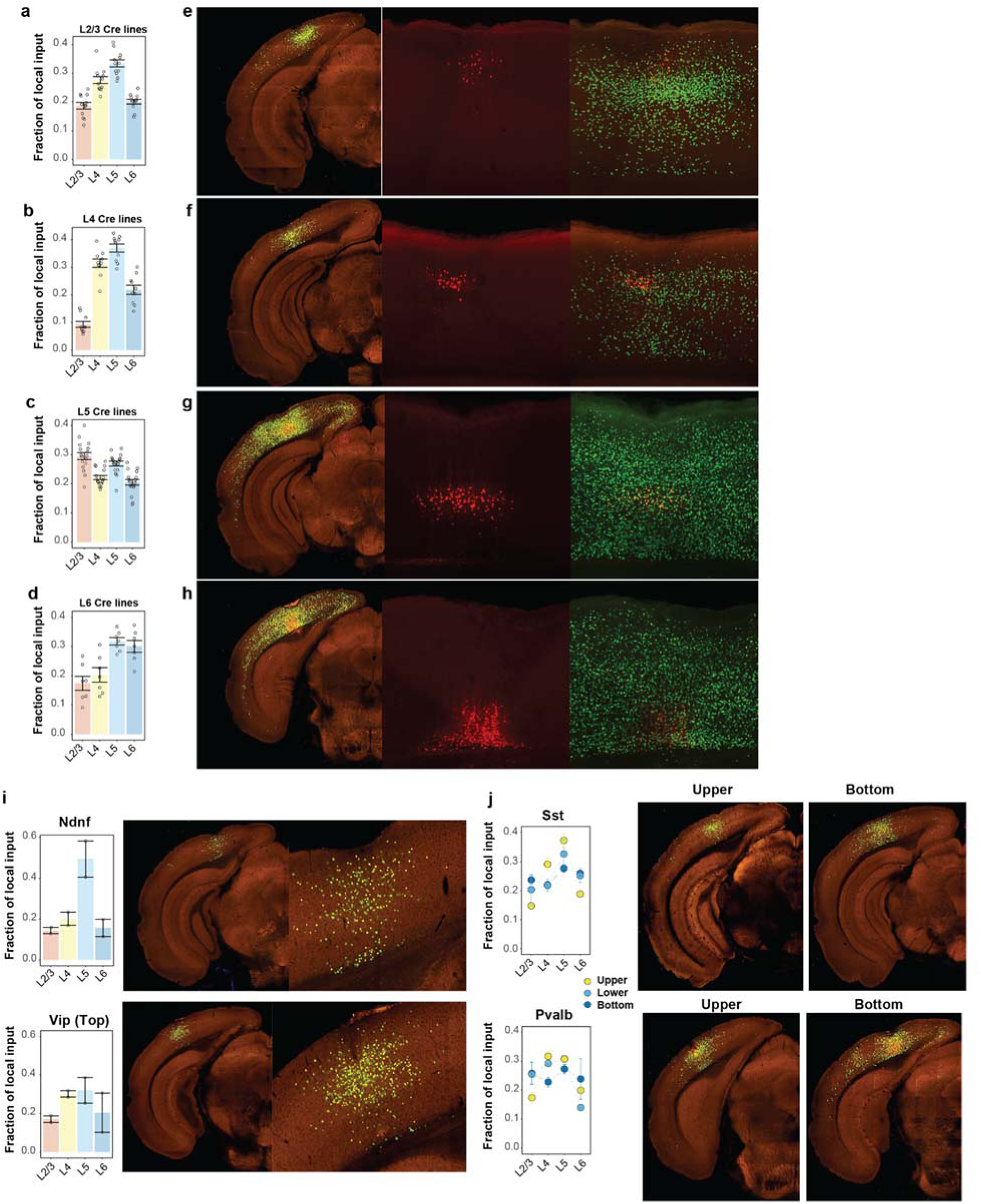
Comparison of local inputs to excitatory neurons and interneurons in different depths of the VISp. **(a-d)** Fraction layer input of ipsilateral VISp inputs to excitatory neurons in L2/3 (a), L4 (b), L5 (c) and L6 (d) of VISp. Data are shown as mean ± s.e.m. **(e-h)** Representative images showing layer-specific local inputs to excitatory neurons in L2/3 (e), L4 (f), L5 (g) and L6 (h) of VISp. The left panels show STPT images of brain sections containing starter cells, and the right two panels are confocal microscopic images showing the distribution of starter cells and local inputs. Starter cells are identified by the coexpression of dTomato from the AAV helper virus and nucleus-localized EGFP from the rabies virus. **(i)** Comparison of local input patterns to Ndnf-Cre and Vip-Cre experiments with Top distribution of starter cells. Data are shown as mean ± s.e.m. **(j)** Comparison of local inputs to Sst and Pvalb experiments with different depths of starter cell distribution. Representative images containing the injection sites are provided for Sst-Cre and Pvalb-Cre experiments in the Upper and Bottom groups. Data are shown as mean ± s.e.m.

Considering the layer-specific local input patterns observed in excitatory neurons, we compared local input patterns of interneurons among the four different groups based on the depth of starter cell distributions (**Figure 6i-j**). The Top group includes two Ndnf experiments and two Vip experiments, with the Ndnf starter cells receiving the most inputs from L5, and the Vip starter cells receiving strong inputs from L4 and L5. The Upper, Lower, and Bottom groups each exhibit distinct local input patterns, with the Bottom group receiving more inputs from L6 than the Upper group. Our results suggest cell-class- and layer-dependent local input patterns for both excitatory neurons and interneurons.

### Hierarchical order of mouse visual areas defined by presynaptic inputs

We subsequently explored whether the laminar distribution of presynaptic inputs to visual areas reveals the hierarchical ordering of these areas. In primates, the hierarchy of visual cortical areas was derived by designating anatomical connections to feedforward and feedback directions, with feedforward connections originating from superficial layers in a lower area and terminating in L4 in a higher area, and feedback connections originating from deeper layers in a higher area and terminating outside L4 in a lower area^58^. Other studies used the fraction of labeled presynaptic supragranular neurons (SLN), defined as the number of labeled neurons in L2/3 divided by the sum of labeled neurons in supra- and infra-granular layers (L2/3 + L5 + L6), to derive a hierarchical ordering of the primate visual cortex areas that was consistent with the Felleman and Van Essen hierarchy^58, 59^. Unlike what was described for primates, L4 neurons in mouse VISp do appear to play an important role as well as other layers in information relay to HVAs. A preference for L4 inputs from VISp is particularly noticeable for VISrl and SSp-bfd-rll, with the highest fraction of VISp inputs coming from L4 (**Supplementary Figure 12**).

In an effort to identify a quantitative hierarchical parameter for visual cortex areas using retrograde labeled cells, we first compared the laminar distribution of presynaptic inputs for the connections between VISp and VISam (**Figure 7a**). In our previous cell-class-specific axon projection mapping^49^, we predicted that VISp lays at the base of the visual area hierarchy, whereas VISam resides at the top of the hierarchy. Accordingly, inputs from VISam to VISp are considered feedback inputs, whereas those from VISp to VISam are considered feedforward inputs. We find that the percentages of L2/3 and L4 inputs of the VISp-to-VISam connection are significantly higher than those of the VISam-to-VISp connection (*P* < 0.001, two sample t-test), and the percentage of L5 inputs from VISam to VISp is significantly higher than that from VISp to VISam (*P* < 0.001, two sample t-test). In contrast, the percentage of L6 input does not show statistically significant difference between the two directions (*P* > 0.05, two sample t-test). Examination of visual area inputs to VISp and VISam further confirmed that the fractions of input from L2/3/4 and L5 present a complementary pattern consistent with the predicted relative hierarchical positions of each source area and target area based on laminar projection patterns (**Figure 7b-c**). We also compared the laminar distribution of inputs from various ipsilateral cortical areas to VISp (**Figures 7d-f**), and find that inputs from almost all other cortical areas to VISp show lower fraction of L2/3/4 inputs and higher fraction of L5 inputs than intra-VISp inputs, consistent with separate roles of L2/3/4 and L5 in feedforward and feedback information relay. In contrast, the fraction of L6 input shows a lateral to medial gradient with fraction of L6 input higher in the lateral areas and lower in the medial area.

**Figure 7.**
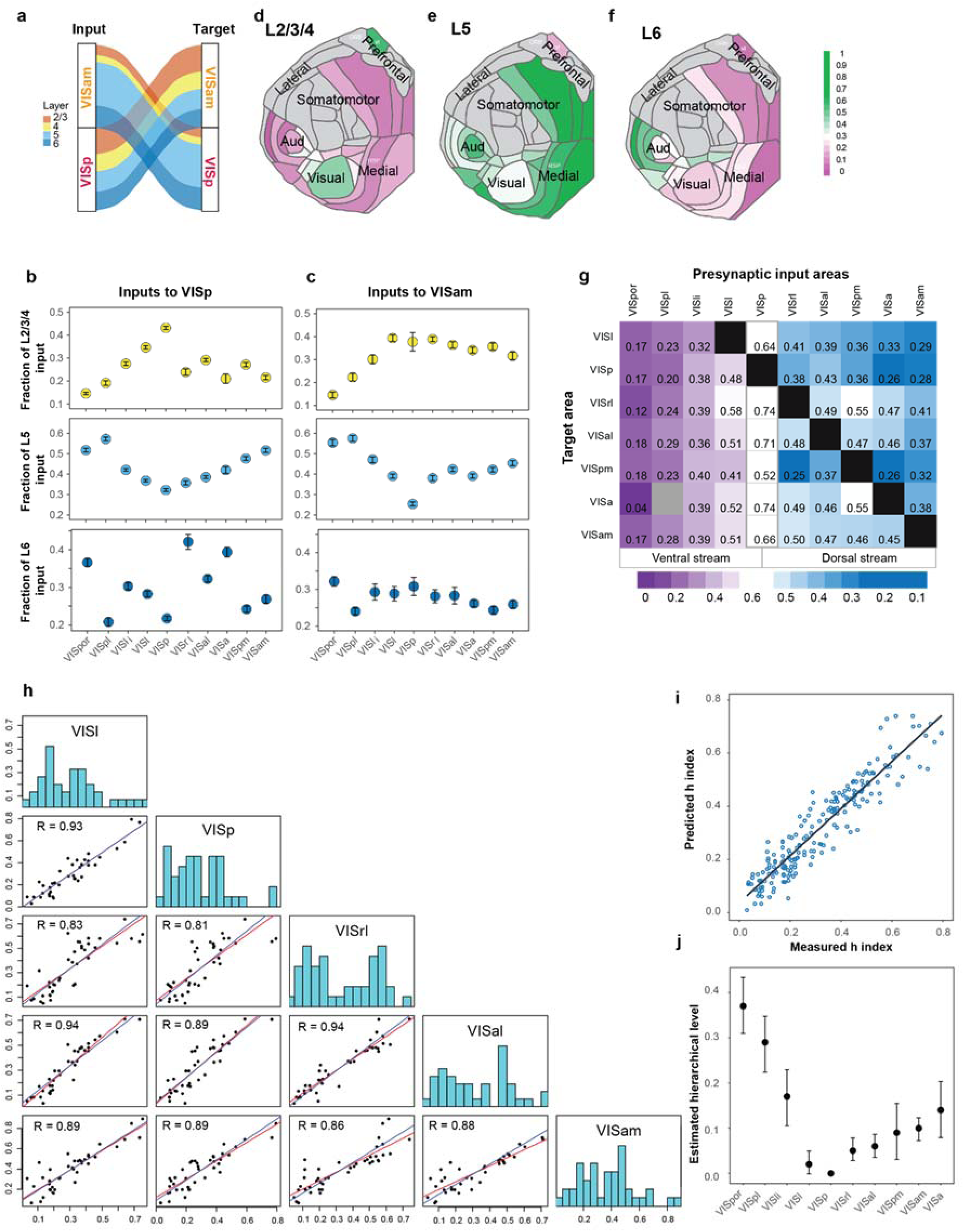
Relative hierarchical positions of the primary visual cortex and higher visual areas. **(a)** Laminar distribution of inputs for connections between VISp and VISam. Fractions of total inputs in the source area across layers were calculated for each experiment. Mean fraction of layer-specific inputs was used to represent the laminar distribution of inputs between the source area and the target area. **(b-c)** Comparison of laminar distribution of visual area inputs to VISp (b) and VISam (c). Fractions of total inputs in the source area across layers were calculated for each experiment. Each dot represents the mean (± s.e.m.) fraction of layer-specific input from the source area to the target area. Visual areas are organized based on previously predicted hierarchical positions and separated into the dorsal and ventral streams. **(d-f)** Comparison of the fraction of L2/3/4 input (d), L5 input (e) and L6 (f) input from various cortical areas to VISp. Mean fraction layer input was calculated to represent the laminar distribution of inputs between the source area and VISp. Each source area is colored according to the mean fraction of inputs in L2/3/4 (d), L5 (e) and L6 (f), respectively. **(g)** Matrix of *h* index of inputs from the 10 visual areas to the 7 targets. *h* index is calculated as the ratio of inputs in layers 2/3/4 to the sum of inputs in layers 2/3/4/5. The visual areas are separated into the ventral and dorsal stream. Each cell within the matrix represents the mean *h* index value of inputs in a given source area to a target area. Gray cell represents no availability of data due to sparse inputs from the source area. **(h)** Pairs plots showing the correlation of measured *h* index values of cortical source areas sending inputs to specific pairs of target areas. Each point represents the average pair of *h* index values obtained in a given source area to a pair of target areas as indicated at the top and the right of each graph. VISa is not included as one of the target areas due to a low number of experiments in VISa, and VISpm is excluded due to the lack of experiments in L4. The red lines are the best fit lines (least-squares regression lines), and the blue lines are the lines with a slope equal to 1 that best fit the points. **(i)** Correlation between measured and predicted *h* index values between cortical source areas and the five target visual areas in panel h. We used the linear regression analysis to estimate a set of hierarchical levels that best predict the measured *h* values. A model can be specified as Y = Xβ, wherein Y is a vector containing the *h* values of all source areas to each target, β contains the estimated hierarchical levels assigned to each area, and X is the incidence matrix. X is constructed so that each column corresponds to one of the 43 cortical areas and each row corresponds to a connection between two areas. All of the elements of a row are zero except in the two columns corresponding to the areas participating in the connection, with the source area taking the value of −1 and the target area taking the value of 1. The hierarchical level of VISp was set at zero. **(j)** Estimated hierarchical levels obtained by the linear regression model. The hierarchical level of VISp was set at zero. Error bars indicate 90% confidence intervals. Visual areas are separated into the dorsal and ventral streams (to the right and left of VISp, respectively).

Our observations suggest that the ratio of inputs from the superficial L2/3/4 to the sum of inputs from the superficial layers and L5 (hereinafter referred to as the *h* index) could be used to quantify the hierarchical positions of the mouse visual areas. We calculated the average *h* index for presynaptic inputs from 10 source visual areas to 7 target visual areas (**Figure 7g**). For a given target, inputs from VISp exhibit the highest *h* index values as compared to the HVAs, consistent with VISp at the lowest hierarchical position, and inputs from HVAs of higher hierarchical positions in the ventral and dorsal streams have lower *h* index values as compared to other HVAs in the same stream.

To explore whether *h* index can serve as a quantitative parameter for hierarchical distance between cortical areas, we performed correlation analysis of *h* values measured between common cortical source areas and paired visual targets (**Figure 7h**). We hypothesized that if the measured *h* index faithfully reflects the hierarchical distance between the target area and the source area, the difference between *h* values measured for a common source area and paired target areas would be the hierarchical distance between the two target areas. This relationship would be translated into a best-fit line with a slope equal to 1 and an intercept indicating the hierarchical distance between the two target areas when plotting paired *h* values measured for common cortical source areas. We compared the paired *h* values between cortical source areas and five target visual areas, with VISpm excluded for lack of L4 experiments and VISa excluded for low sample size. Overall, we found a fair correspondence between the best fit lines based on the least-squares criterion and the best fit with a slope equal to 1. We then fit a linear regression model to the measured *h* values between cortical source areas and the five visual areas with the hierarchical level of VISp set at zero. The relative hierarchical orders of cortical areas were estimated to best predict the measured *h* index values. A strong correlation (R^2^ = 0.94) was found between the predicted *h* values and the measured values (**Figure 7i**). The estimated hierarchical levels for the visual areas (**Figure 7j**) are overall consistent with the predicted hierarchy based on the cell-type-specific projection^49^, with the exception that the linear model fitting *h*-index values places VISa at the top of the dorsal stream instead of VISam. Consistent with our previous findings, we find that the hierarchy in the mouse visual cortex is shallow, especially for the dorsal stream.

## Discussion

Here we present the construction and validation of a retrograde connectome pipeline for the mouse brain, with a focus on the visual cortical areas. With improved virus tools and informatic processing, our pipeline can be utilized to conduct large-scale systematic mapping of brain-wide presynaptic inputs at the cell class or type level. Together with our anterograde projection mapping pipeline, the current work proves the feasibility to build a comprehensive, directional, and 3D connectional atlas of the mouse brain at the cell type level.

### Target, layer and cell class together determine the presynaptic input patterns

Our retrograde connectome dataset reveals that the presynaptic inputs to defined neuronal classes are determined predominantly by the target area, followed by layer distribution of starter cells and Cre line-defined cell classes. Both VISp and HVAs receive the most inputs from the isocortex, followed by the thalamus and HPF. Strong cortical inputs are often from source areas that receive strong visual area inputs, indicating reciprocal connections between the visual cortex and other cortical modules. Each target area in the visual cortex exhibits unique input patterns, distinguishing the dorsal stream targets from the ventral stream targets, and the anterolateral HVAs from the anteromedial HVAs. In contrast to the highly differential cell-class-specific anterograde projection patterns^49^, rabies tracings from Cre-defined cell classes in the same target reveal overall similar global input patterns. We have obtained similar results when we applied the same approach described in this study to another mouse cortical area, the primary motor cortex^78^. Nonetheless, we also observe quantitative differences in inputs to layer- and Cre-defined starter cell classes. Consistent with the feedforward thalamocortical connectivity from LGd to VISp, we find that L4 excitatory neurons receive the highest LGd input and lowest higher-order brain area inputs as compared to excitatory neurons in other layers. We also discover that L6 neurons receive more contralateral cortical inputs than any other layers, despite that the intracortical projections of L6 neurons are often locally restricted. Among the interneurons, we also find that inputs to interneurons are mainly determined by target area and layer distribution of starter cells, although cell-class-dependent local and long-range inputs can also be observed. Our observation is in line with a recent rabies based tracing study which found strong similarity in local and long-range inputs to L2/3 excitatory and inhibitory neuron types and significantly lower fraction of contralateral inputs to these L2/3 neurons as compared to L6 Ntsr1 neurons^79^.

### Layer 4 neurons in visual cortex send significant inputs to other cortical areas

Our study reveals that the role of L4 excitatory neurons is not limited to being the receiver of feedforward information. Inputs from L4 and L2/3 together contribute to the feedforward interareal connections. In particular, VISp inputs to SSp-bfd-rll are almost exclusively from L4, and similar preference for L4 inputs is observed for the connection between other HVAs and SSp-bfd-rll. For HVA inputs to VISp, most of the input cells are in deep layers of HVAs, consistent with the classical dogma of feedback information flow, while substantial amounts of inputs reside in L4 of HVAs, suggesting a broad role of L4 neurons in the processing of visual information. Being the major subclass of neurons receiving thalamic inputs while also sending outputs to HVAs, VISp L4 neurons could in principle facilitate direct integrated processing of visual information. Recent studies also found that L4 neurons send long distance projections and are the major source of visual cortex projections to Pvalb neurons in barrel field^40^, consistent with a more complex role of L4 excitatory neurons in interareal connectivity.

### Distinct patterns of thalamic projections to the visual areas

The thalamus sends the second most abundant inputs to the visual areas. VISp is distinct from the HVAs by preferentially receiving inputs from LGd as compared to LP, and L4 of the VISp receives the most abundant LGd inputs as compared to other layers, consistent with the feedforward flow of visual information from the LGd to L4 of the VISp. The overall patterns of thalamic inputs to the HVAs show clear correlation with the anatomical proximity of HVAs. VISrl and SSp-bfd-rll are either bordering or located in barrel cortex, and both receive abundant inputs from the VPM, reminiscent of the strong VPM input to neurons in barrel cortex^40, 80^. The SSp-bfd-rll also receives substantial projections from anterior VISp but barely from posterior VISp^49, 50^. In this study, we find that SS-bfd-rll sends projections back to visual areas, demonstrating reciprocal connections between them^81^. Unlike other HVAs that receive the highest proportion of thalamic inputs from LP, SSp-bfd-rll distinctively receives the most inputs from PO and VPM, suggesting that SSp-bfd-rll could be a transition area integrating the processing of both visual and somatosensory information.

### Hierarchical organization of the mouse visual cortex

We explore the possibility of predicting the hierarchical positions of mouse visual cortical areas by the laminar distribution patterns of presynaptic neurons, an alternative approach from predictions based on laminar termination patterns of axon projections^49^. The feedback and feedforward connections in the mouse visual areas are gauged by the ratio of L2/3/4 inputs to L2/3/4/5 inputs. Using this quantitative parameter, we obtain a shallow hierarchy of the visual areas, which places VISp at the bottom, and VISa and VISpor at the top of the hierarchy in the dorsal stream and the ventral stream, respectively. This hierarchy is largely consistent with the one we previously derived based on axon termination patterns^49^, and the anatomical hierarchy revealed by both output and input connectivity patterns mirrors the functional hierarchical organization of mouse cortical visual areas^82^. Our retrograde tracing shows VISa at the top of dorsal stream hierarchy, which is different from anterograde tracing showing VISam at the top. In primates, temporal association cortex (TE) in the ventral visual cortical pathway has large overlapping visual receptive field with no clearly separated visuotopic map^66, 83^. Similarly, VISa in mouse dorsal stream has larger receptive field than VISam^53^, and VISa doesn’t have a complete visual field, whereas VISam does^54, 55^. Our current study also demonstrates that VISa receives more input from SSp than VISam does (**Figure 3c**). These differences between VISa and VISam support our current finding that VISa is higher in hierarchical level than VISam. Since we used cell-class specific Cre lines to define the starter cells in retrograde tracing, a comprehensive coverage of cell types in the target is a prerequisite for defining the hierarchical orders using presynaptic input patterns. Compared to our previous larger-scale anterograde projection study covering nearly the entire corticothalamic system^49^, our current study is restricted to the visual cortex and has varying levels of coverage for different visual areas. As we continue our effort to build the Allen Mouse Brain Connectivity Atlas, the brain-wide cellular-level retrograde connectome will enable a more in-depth understanding of the organizing principles of the brain.

### Limitations of rabies virus tracing

The monosynaptic rabies virus tracing system is a powerful tool in its ability to selectively infect starter cells and label only the first-order presynaptic neurons. However, although we have improved our virus tools to further enhance specificity and efficiency, there are still limitations of this strategy, which should be taken into consideration when interpreting the results.

Due to the high affinity between TVA and EnvA, low-level leaky expression of TVA in the absence of Cre is sufficient for rabies infection^29, 32, 34, 61^. Although the leaky expression of RG is often too low to allow trans-synaptic transportation of rabies virus, these cells can be mistakenly counted as local trans-synaptically labeled cells. Our AAV helper virus is specifically designed to reduce spurious expression in the absence of Cre by utilizing TVA^66T^. However, many of the Cre-driver lines also have expression in areas sending input to the visual area, and AAV1 serotype is known for its ability to retrogradely label the soma by traveling along the axon. It is possible that the AAV helper virus can infect neurons in the brain area with direct input to the visual area, in effect creating new starter cells, leading to the labeling of neurons in areas without direct connection with the visual area. Therefore, independent connection mapping strategies are required to verify novel connections revealed by rabies tracing.

On the other hand, it is also possible that the rabies virus tracing system does not reveal all presynaptic neurons, even though we used the CVS-N2c strain which has higher trans-synaptic efficiency. Rabies virus may not cross all synapses with equal efficiency, leading to preferential representation of certain cell types within the presynaptic connectome. The efficiency of the trans-synaptic spread of rabies virus is also affected by the expression level of RG in the starter neurons, which is in turn limited by the expression of Cre from the driver lines, the titers of the AAV helper virus and rabies virus tracer, as well as the amount of virus particles successfully delivered to the target sites. The potential incompleteness and bias of the retrograde connectome mapped by the rabies tracing system should always be considered, especially when an understanding of the absolute number and the strength of synaptic connections between the input cells and the starter cells is desired.

## Author contributions

H.Z., J.A.H., A.C., S.M. and S.Y. contributed to overall project design. A.C. designed and orchestrated the viral tracing technology as well as viral production capability and established these with the help from S.Y. T.Z. and M.T.M. S.Y., T.Z. and M.T.M. performed virus production. A.C., S.Y., T.L.D. and B.O. conducted initial proof-of-principle studies. A.W. and P.A.G. supervised surgical procedures with contributions from B.O., C.N., K.M., S.L, A.C., L.C., K.N., N.H., E.G., J.L., R.A, R.H., and J.S. P.A.G. supervised ISI procedures with contributions from S.C., S.S., E.K.L., F.G., and T.N. M. McGraw. supervised histological processing with contributions from T.E., J.B., M. Maxwell., H.G., A.G., K.B., and A.R. P.R.N. coordinated imaging procedures with contributions from R.E., M.G., S.R., L.P., N.I.D., N.K.N., and M.J.T. L.N., L.K. and W.W. performed informatics data processing. K.E.H. and S.Y. coordinated workflow and carried out quality control. M.N. contributed to the development of data visualization tools. Q.W., S.M., J.A.H., A.C., S.Y. and H.Z. formulated data generation and analysis strategies. S.Y. analyzed data and prepared figures. S.Y., B.T., and H.Z. wrote the manuscript with inputs from all authors.

## Acknowledgment

We are grateful to the Transgenic Colony Management, Neurosurgery & Behavior, Lab Animal Services, Molecular Genetics, Imaging, Histology, Technology, and Project Management teams at the Allen Institute for technical and management support. We thank Thomas R Reardon, Andrew J Murray and Ian Wickersham for providing cell lines and plasmids for the establishment of rabies virus production at the Allen Institute. This work was supported by the Allen Institute for Brain Science and by the National Institute of Mental Health (NIMH) of the National Institutes of Health (NIH) under award number U19MH114830 to H.Z. The content is solely the responsibility of the authors and does not necessarily represent the official views of NIH and its subsidiary institutes. We thank the Allen Institute founder, Paul G. Allen, for his vision, encouragement, and support.

## METHODS

All experimental procedures related to the use of mice were approved by the Institutional Animal Care and Use Committee of the Allen Institute for Brain Science, in accordance with NIH guidelines.

### Outline of the data generation and processing pipeline

A standardized data generation and processing platform was established. Mice first received virus injection, and the brains were then imaged by serial two-photon tomography (STPT). Images passed annotation quality control (QC) were subject to data processing through the informatics pipeline, while brain sections around the injection sites were mounted, immunostained to enhance the red fluorophore and imaged by confocal microscopy to identify starter cells. Specimens failed the staining QC and starter cell QC were removed from the pipeline, and finally, artifacts in informatics processing were identified and corrected through segmentation QC.

### Data Availability

Plasmids for the generation of recombinant viruses will be deposited in Addgene. All anterograde tracing data (including high resolution STPT images, and informatically processed axonal projection across brain structures) are available through the Allen Mouse Brain Connectivity Atlas portal (http://connectivity.brain-map.org/). A link for each anterograde tracing experiment is provided in Supplementary Tables 4 and 5. Original images for transsynaptic rabies viral tracing will be available through the Brain Image Library (https://www.brainimagelibrary.org/). Normalized presynaptic input volumes across the brain for all rabies virus tracing experiments are listed in Supplementary Table 3. Original retrograde labeling data from initial informatic quantification are available upon request.

### Animals

To identify presynaptic inputs to different neuronal populations in the visual cortex, 14 Cre-transgenic mice with distinct cell type and layer labeling patterns (aged 2-6 months, either gender depending on availability) were used. These Cre lines have been previously utilized together with anterograde AAV viral tracers for the construction of the Allen Mouse Brain Connectivity Atlas. The expression patterns of these lines can be found in the Allen Institute Transgenic Characterization data portal (http://connectivity.brain-map.org/transgenic). To quantify the spurious rabies virus labeling, wild-type animals were used, and to compare different recombinant rabies and AAV helper viruses, Cre transgenic mice labeling neurons and non-neuronal cells were used. The specific genotypes used for each experiment are listed in Supplementary Tables 1 and 2. Mice were housed under 14h:10h light-dark cycle with ad libitum access to food and water.

### Virus design, preparation and titer information

AAV viruses and rabies viruses used in the mesoscale retrograde connectome pipeline were generated in the Allen Institute of Brain Science. Our AAV helper viruses utilize the FELX strategy and contain the tricistronic cassettes of Syn-DIO-TVA-dTomato-RV G, followed by a short bovine growth hormone polyadenylation sequence. The Kozak sequences and the starter codon of TVA are located 5’ to the FLEX switch, while the TVA (lacking a start codon)-P2A-dTom-P2A-RV G cassette is within the FLEX cassette and inverted respective to the promoter. The AAV helper viruses selected for the mesoscale retrograde connectome pipeline utilized a mutant TVA, TVA^66T^, and the RV G from the CVS N2c strain.

The AAV1 serotype of the helper virus was produced using a helper-free HEK293 cell system followed by iodixanol gradient purification. Multiple batches of AAV1-Syn-DIO-TVA^66T^-dTom-CVS N2cG viruses were used in the course of the mesoscale connectivity project, and the titers of the viruses were in the range of 2×10^12^ to 1×10^13^GC/ml.

The CVS N2c^dG^-H2B-EGFP rabies virus was generated by replacing GFP in the rabies genomic plasmid RabV CVS-N2c(deltaG)-EGFP (Addgene, Plasmid #73461) with H2B-EGFP flanked by 5’ XmaI and 3’ NheI-KasI sites. EnvA CVS N2c^dG^-H2B-EGFP rabies viruses were generated from the genomic plasmid as described previously^1^. The titer of EnvA CVS N2c^dG^-H2B-EGFP rabies virus used in the study was adjusted to be around 5×10^9^ GC/ml.

The SAD B19^dG^-H2B-EGFP virus was generated by inserting the H2B-EGFP sequence into the gG locus of the pSADdeltaG-F3 plasmid (Addgene, Plasmid #32634). The EnvA SAD B19^dG^-H2B-EGFP rabies viruses were generated from the genomic plasmid as described previously^2^. The EnvA SAD B19^dG^-GFP was from Salk Institute. Both EnvA SAD B19^dG^-H2B-EGFP and EnvA SAD B19^dG^-GFP viruses were diluted to 5×10^9^ GC/ml to match that of the EnvA CVS N2c^dG^-H2B-EGFP virus.

### Surgery

All mice received unilateral injection into a single target region in the left hemisphere. For monosynaptic retrograde tracing of whole brain inputs to Cre-defined cell populations, the AAV helper virus was injected first into the target site, followed 21±3 days later by another injection in the same location with the EnvA CVS N2c^dG^-H2B-EGFP rabies virus. After one week survival, animals were sacrificed, and perfused with 4% paraformaldehyde (PFA). Brains were dissected and post-fixed in 4% PFA at room temperature for 3–6 h and then overnight at 4°C.

To precisely target each visual area, functional mapping of visual field space by intrinsic signal imaging (ISI) was used to guide injection placement^3^. An image of the surface vasculature was acquired to provide fiduciary marker references on the surface of the brain. An overlay of the visual field map over the vasculature fiducials was used to identify the target injection site. For injections that failed repeatedly under the guidance of ISI, transcranial injections were conducted using stereotaxic injection coordinates specific for each target site. The anterior/posterior (AP) coordinates are referenced from the transverse sinus (TS), the medial/lateral (ML) coordinates are distance from midline at Bregma, and the dorsal/ventral (DV) coordinates are measured from the pial surface of the brain. Stereotaxic coordinates for each area are as follows: VISp (6 subareas) (VISp-1: AP:1.50(TS), ML:-2.55, and DV:0.3, 0.6; VISp-2: AP:2.59(TS), ML:-2.55, and DV:0.3, 0.6; VISp-3: AP:1.90(TS), ML:-3.10, and DV:0.3, 0.6; VISp-4: AP:1.05(TS), ML:-3.50, and DV:0.3, 0.6; VISp-5: AP:0.75(TS), ML:-3.00, and DV:0.3, 0.6; VISp-6: AP:0.61(TS), ML:-2.10, and DV:0.3, 0.6), VISl (AP:1.4(TS), ML: −4.10, DV:0.3, 0.6); VISpm (AP:1.9(TS), ML: −1.60; DV:0.3, 0.6), VISam (AP: 3.0(TS), ML: −1.70, DV:0.3, 0.6), VISal (AP: 2.4(TS), ML: −3.70, DV:0.3, 0.6, Angle:15°), and VISrl (AP: 2.8(TS), ML:-3.30, DV: 0.3, 0.6). For some target areas, injections were made at two depths to label neurons throughout all six cortical layers. The AAV1 helper virus was injected using the iontophoresis method, with current settings of 3 µA, 7 sec on/off cycles and 5 min total. The EnvA rabies viruses were injected using a nanoinjector, and a total of 500 nl was delivered in 23 nl increments over a 3 min and 10 sec interval.

Tamoxifen-inducible Cre line (CreER) mice were treated with 0.2 mg/g body weight of tamoxifen solution in corn oil via oral gavage once per day for 5 consecutive days starting the week following virus injection. Trimethoprim-inducible Cre line mice were treated with 0.3 mg/g body weight of trimethoprim solution in 10% DMSO via oral gavage once per day for 3 consecutive days starting the week following AAV virus injection. Rabies viruses were injected 3 weeks post induction. All mice were deeply anesthetized before intracardial perfusion, brain dissection, and tissue preparation for serial imaging.

### STPT

The injected brains were imaged by STPT (TissueCyte 1000, TissueVision Inc. Somerville, MA) as described previously with a few modifications^3,4^. In brief, brains were embedded in agarose block, and imaged from the caudal end along the rostrocaudal z-axis. The specimen was illuminated with 925 nm wavelength light. Two-photon image tiles for red and green channels with a nominal resolution of 0.875 μm x 0.875 μm x 2 µm x-y-z were taken at 40 μm below the cutting surface. The laser power and photo-multiplier tube (PMT) voltage were set at 190 mW (measured at the objective) and ∼600 V (equal on all channels). In order to compensate for variation between imaging systems and specimens, these parameters are adjusted on each imaging run using an observed level of autofluorescence in the red channel. The following procedures were conducted: locate the central canal of the brain stem; locate the surface of the tissue; adjust the objective piezo stage such that the image plane is 40 μm deep in the specimen; move 700 μm laterally, exposing an area of uniform tissue structure; adjust the PMT voltage such that the mean intensity of this area falls within the range of 600-650; from the central canal, the specimen is then centered laterally within the imaging area and the acquisition is commenced. After an entire brain section was imaged, a 100-μm section was removed from the specimen by the vibratome, followed by imaging of the next plane. Scanned image tiles were stitched to form a single high-resolution image. Images from 140 sections were collected to cover the full range of mouse brain. Upon completion of imaging, sections were retrieved and stored in PBS with 0.1% sodium azide at 4°C.

### Starter cell identification and quantification

The starter cells are those with both AAV helper virus and EnvA RV-H2B-EGFP infection, and thus have red fluorescence in the soma and green fluorescence in the nuclei. For starter cell quantification, TissueCyte brain sections were sorted according to the rostrocaudal axis. Around 20 100-μm sections flanking the virus injection sites were identified, mounted on gelatin coated glass slides, and immunostained to enhance red fluorescence signal. The immunofluorescence staining was conducted using an automated slide stainer (Biocare, IntelliPATH FLX). Slides were blocked in Image iT FX Signal Enhancer (Thermo Fisher Scientific Cat# I3693) for 45 minutes, followed by 1-hour incubation in a blocking solution containing 1% normal goat serum (Vector Laboratories Cat#S1000) and 1% Triton X (VWR). Sections were then incubated in the primary antibody solution (1% goat serum, 1% Triton X, Rockland Cat# 600-401-379, RRID:AB_2209751, 1:2000) for 1.5 hours, and then the in secondary antibody solution (1% goat serum, 1% Triton X, and 1:500 goat anti-rabbit conjugated with Alexa Fluor 594, Thermo Fisher Scientific Cat# A-11037, RRID:AB_2534095) for 2 hours at room temperature after rinsing with 0.1% Triton X wash solution. All sections were stained with 5 μM Dapi (Thermo Fisher Scientific D1306) and coverslipped using Fluoromount G (Southern Biotech Cat# 0100-01B). Stained sections were imaged using a Leica SP8 TCS confocal microscope under a 10x objective. Starter cells were counted in ImageJ using the Cell Counter plugin.

### Image data QC and annotation

The acquired TissueCyte images and the confocal images went through several steps of quality control processes: annotation QC, staining QC, starter cell QC, and segmentation QC. Specimens that did not pass any one of the QC steps were considered fails and removed from the pipeline. After TissueCyte imaging, specimens are assessed for surgical and imaging quality through Annotation QC. Failures at this step include no green signal, TissueCyte imaging error, tissue damages, and poor surgical targeting. Polygons are drawn around the injection site to link the injection site to the Allen Mouse Brain Common Coordinate Framework, version 3 (CCFv3). Specimens passed Annotation QC were sent for the next step of mounting, and immunostaining for starter cell identification. Staining QC identified and removed specimens in which no red-fluorescent cells were found after immunostaining-mediated enhancement of red fluorescence signal from the AAV helper virus. Starter cell QC further removed specimens in which no starter cells were identified after confocal imaging. Finally, specimens with errors in the subsequent informatics data pipeline steps were identified in the Segmentation QC step.

### Image data processing

Images were processed and registered to the CCFv3 through our informatics data pipeline (IDP)^5,6^. The signal detection algorithm was modified to detect nuclear objects with high sensitivity, which accepts out of focus nuclei and has lower contrast requirements. In addition, high intensity pixels near the detected objects were included into the signal pixel set. Detected objects near hyper-intense artifacts occurring in multiple channels were removed. The output is a full resolution mask that classifies each pixel as either signal or background. An isotropic 3D summary of each brain is constructed by dividing each image series into 10 µm × 10 µm × 10 µm grid voxels. Total signal is computed for each voxel by summing the number of signal-positive pixels in that voxel. Each image stack is registered in a multi-step process using both global affine and local deformable registration to the 3D Allen mouse brain reference atlas as previously described^5,6^.

### Analysis of whole brain presynaptic inputs to the visual areas

The accuracy of targeting was verified by overlaying the injection site polygon of each experiment to the ISI image or by identifying the anatomical structure where the injection centroid was located in the CCFv3. Since the signal detection algorithm was optimized to detect sparse presynaptic labeling with high sensitivity, the automatically detected volume of input signal can have false positives where high background signal is falsely identified as input signal. False positives tend to occur more frequently in brain structures with low input signal and high background fluorescence such as the cerebellum, and are rarely found in areas with strong input signals such as the isocortex. In order to remove this type of artifacts, we identified a set of 92 negative brains that were processed through the pipeline, but showed no rabies-mediated GFP expression, and used this negative dataset to calculate the threshold of false positive signal, i.e., the value of mean input signal volume plus 6 standard deviations for each of the 314 ipsilateral and 314 contralateral major structures of the brain. Any structure not passing this threshold was set to “0”. A manually validated binary mask was then applied to further remove artifacts in informatically-derived measures. Following these two steps, input signal volume in a given structure was normalized to the total input of the brain. The post-threshold, masked, normalized input signal volumes were used to build the weighted connectivity matrix.

When analyzing the fraction of inputs in a given cortical area across layers, the threshold for per structure input signal volume was set at 0.0004. Any structure below this threshold was set as “0”, and no fraction of layer inputs was calculated. This threshold value is higher than 99% of input signal volumes measured for structures in the negative dataset, and is equivalent to around 10 labeled cells based on our comparison of input signal volume and manual counting. We reasoned that cortical structures below this threshold have very sparse RV-labeled neurons, which could lead to extreme values when calculating the layer-specific contribution of inputs.

Hierarchical clustering in Figure 2 was conducted using the pvclust package in R. The agglomerative method used in hierarchical clustering was “ward.D”, and the distance measure used was correlation. The R software was used for statistical tests and generation of graphs.

### Estimation of hierarchical levels

We first identified a quantitative hierarchical parameter *h* based on the anatomical features of feedback and feedforward connections, with *h* calculated as the ratio of layers 2/3/4 input to layers 2/3/4/5 input. We used the linear regression analysis to estimate a set of hierarchical levels that best predict the measured *h* values. A model can be specified as Y = Xβ, wherein Y is a vector containing the *h* values of all source areas to each target, β contains the estimated hierarchical levels assigned to each area, and X is the incidence matrix. X is constructed so that each column corresponds to one of the 43 cortical areas and each row corresponds to a connection between two areas. All of the elements of a row are zero except in the two columns corresponding to the areas participating in the connection, with the source area taking the value of −1 and the target area taking the value of 1. The hierarchical level of the primary visual area was set at zero.

---

**Supplementary Figure 1.**
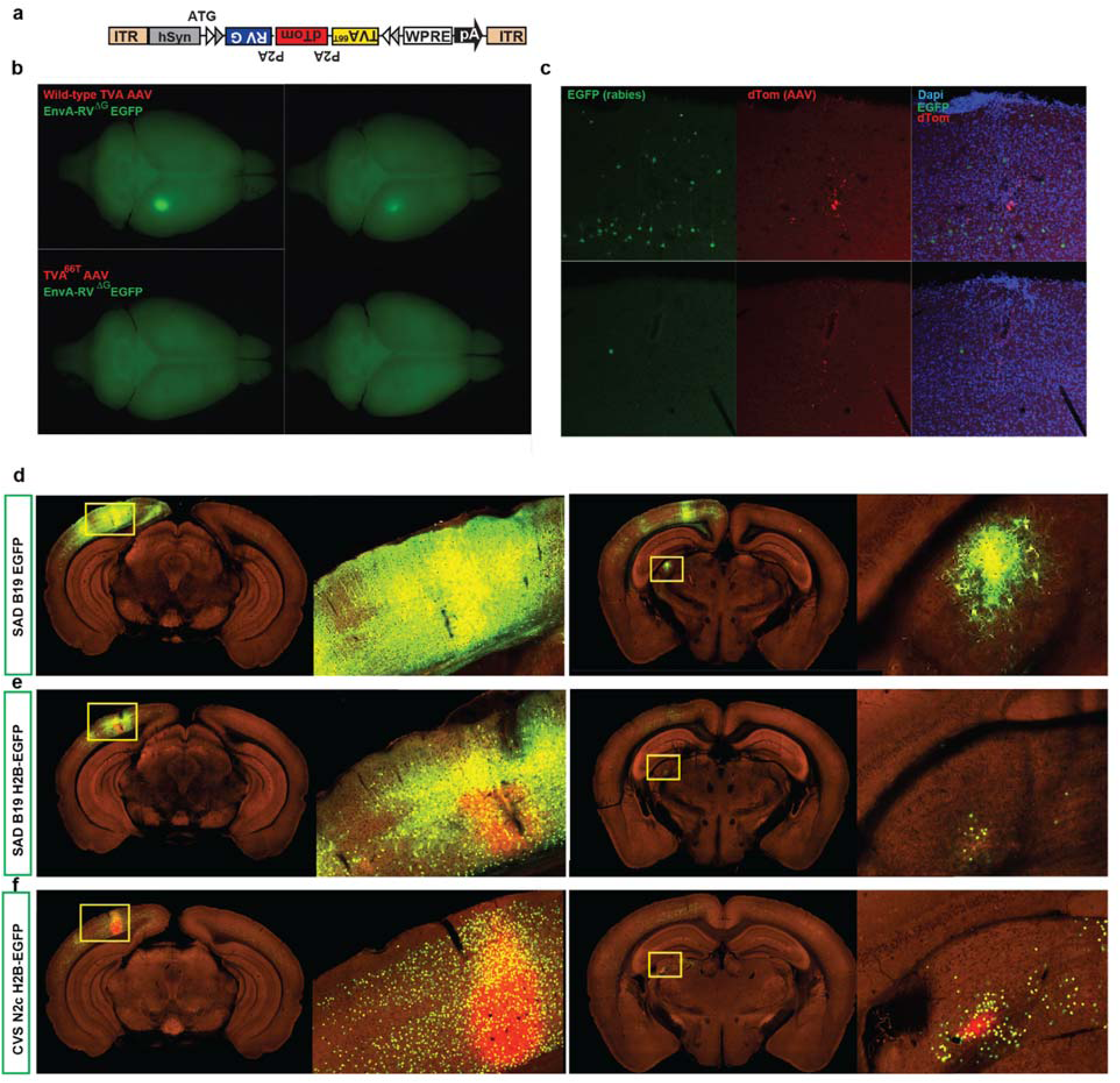
Comparison of different AAV helper viruses and rabies viruses for monosynaptic retrograde tracing. **(a-c)** Comparison of spurious rabies infection from AAV helper viruses expressing wild-type TVA and the mutant TVA^66T^. Tricistronic AAV helper viruses were constructed to conditionally express either the wild-type TVA or TVA^66T^, together with dTomato and RG (a). Cre negative wild-type mice were sequentially injected with AAV helper viruses and EnvA-pseudotyped recombinant rabies viruses expressing EGFP. Each AAV helper virus/rabies virus pair was tested in two wild-type mice. Top-down view of whole brains (b) and observation of the injection sites under the confocal microscope (c) revealed fewer spurious rabies infection from AAV helper viruses conditionally expressing TVA^66T^, dTomato, and RG and EnvA-pseudotyped recombinant rabies viruses expressing EGFP. **(d-f)** Comparison of monosynaptic retrograde tracing in VISp using SAD B19 strain of recombinant RV expressing EGFP (d) or H2B-EGFP (e) or CVS N2c strain of recombinant RV expressing H2B-EGFP (f).

**Supplementary Figure 2.**
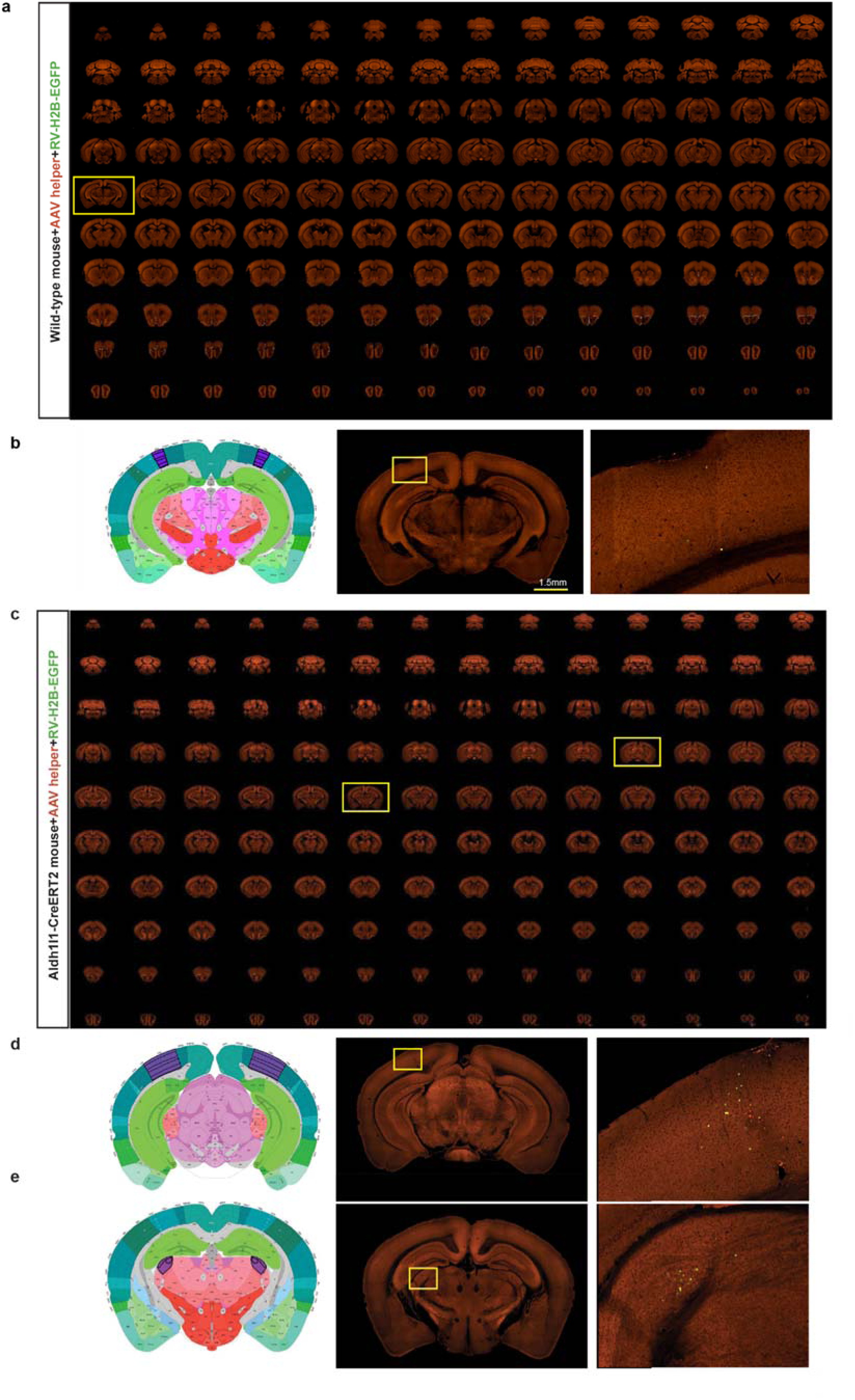
Validation of the AAV helper virus and recombinant rabies used in the mesoscale connectomics pipeline in wild-type mice and non-neuronal Cre lines. **(a)** Sequential two-photon images of a Cre negative wild-type mouse brain injected with the AAV helper virus and EnvA-pseudotyped CVS N2c^dG^ rabies viruses expressing H2B-EGFP. **(b)** Absence of RV-labeled neurons except a few H2B-EGFP-expressing cells in the injection site. Virus injection was targeted to VISp and validation was conducted in two wild-type mice. Left and middle panels: corresponding 2D atlas plate of Allen CCFv3 and image showing the injection site. Right panel: Image magnified from the outlined box in the middle panel. **(c)** Sequential two-photon images of rabies labeling in an astrocyte-specific Cre mouse brain injected with hSyn-driven AAV helper virus and recombinant rabies virus. **(d-e)** Left and middle two panels: corresponding 2D atlas plates of Allen CCFv3 and images showing the injection site. Right panels: Representative images magnified from the outlined boxes in the middle panels reveal the sparse labeling around the injection site (d) and in the LGd (e).

**Supplementary Figure 3.**
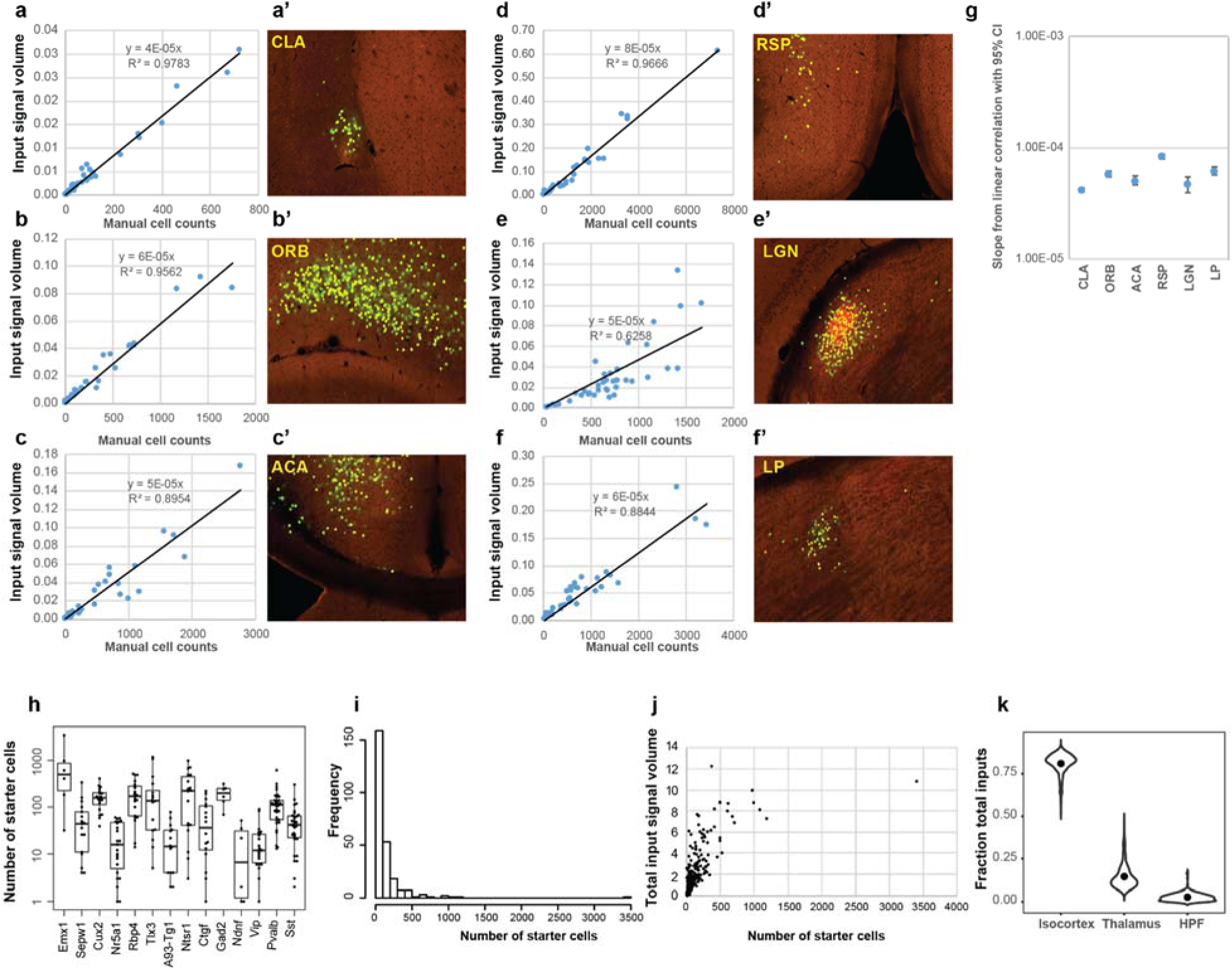
Overview of automatic input signal detection and characterization of inputs to cell classes defined by Cre lines in the visual cortex. **(a-f)** Relationship between per structure input signal volume measured by the informatics data pipeline and cell counts. Linear correlation between input signal volume and manually counted input cells was shown in various brain areas **(a’-f’)**. **(g)** Slopes from linear correlations between informatically measured input signals and manual cell counts in various brain areas. **(h)** Number of starter cells for each experiment categorized in Cre lines. **(i)** Distribution of numbers of starter cells across all experiments. **(j)** Relationship between numbers of starter cells and total inputs from the whole brain. **(k)** Fractions of inputs from isocortex, thalamus and HPF to the mouse visual cortex. Dots represent the median values of input signals.

**Supplementary Figure 4.**
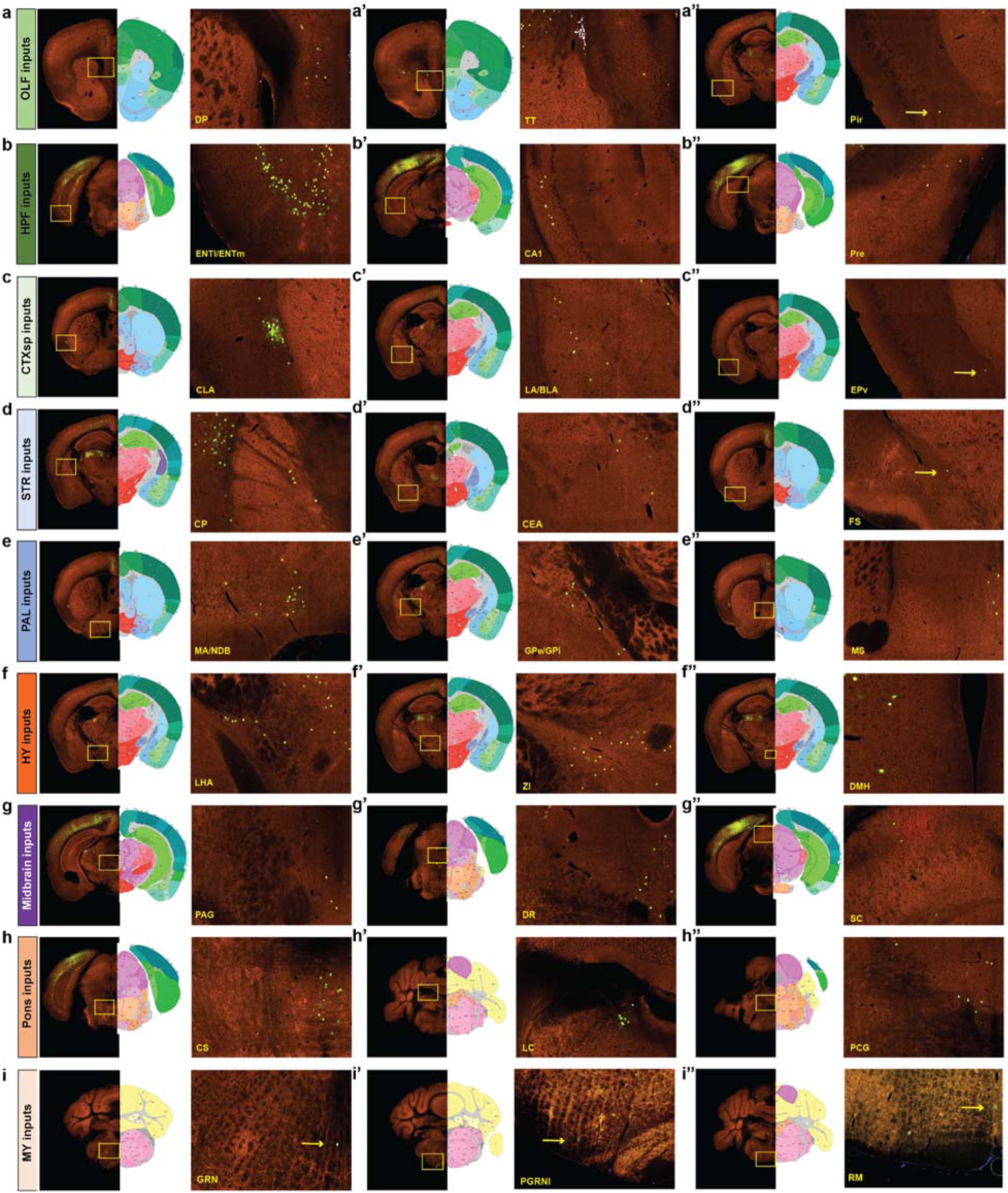
Representative images of presynaptic inputs to the visual areas from anatomical structures outside of the cortex and thalamus. Coronal STPT images and their corresponding 2D atlas plates in Allen CCFv3 show labeled presynaptic neurons in OLF (a-a’’), HPF (b-b’’), CTXsp (c-c’’), STR (d-c’’), PAL (e-e’’), HY (f-f’’), midbrain (g-g’’), Pons (h-h’’), and MY (i-i’’). Enlarged views of boxed areas are shown in the right-hand panels for each major brain region. Arrows highlight the location of single labeled cells.

**Supplementary Figure 5.**
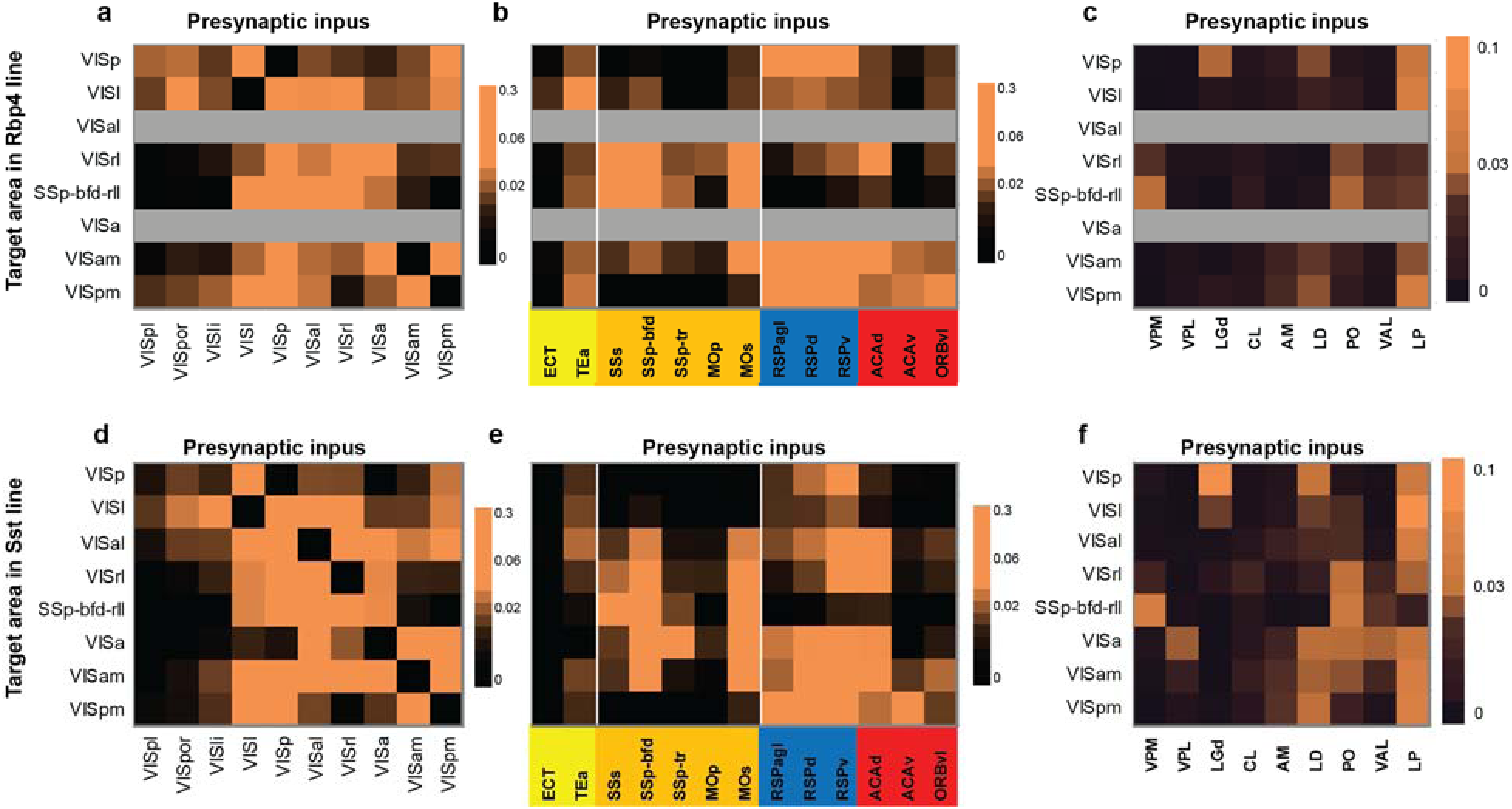
Matrices comparing the inputs to Rbp4 (a-c) and Sst (d-f) neurons in the visual areas from within the visual cortex, non-visual isocortical modules, and thalamus. Gray indicates experiments not available. In the matrix, each row represents experiments with the same target area, and each cell shows the fraction of the total input signal in a given structure measured from a single experiment or the average when n > 1.

**Supplementary Figure 6.**
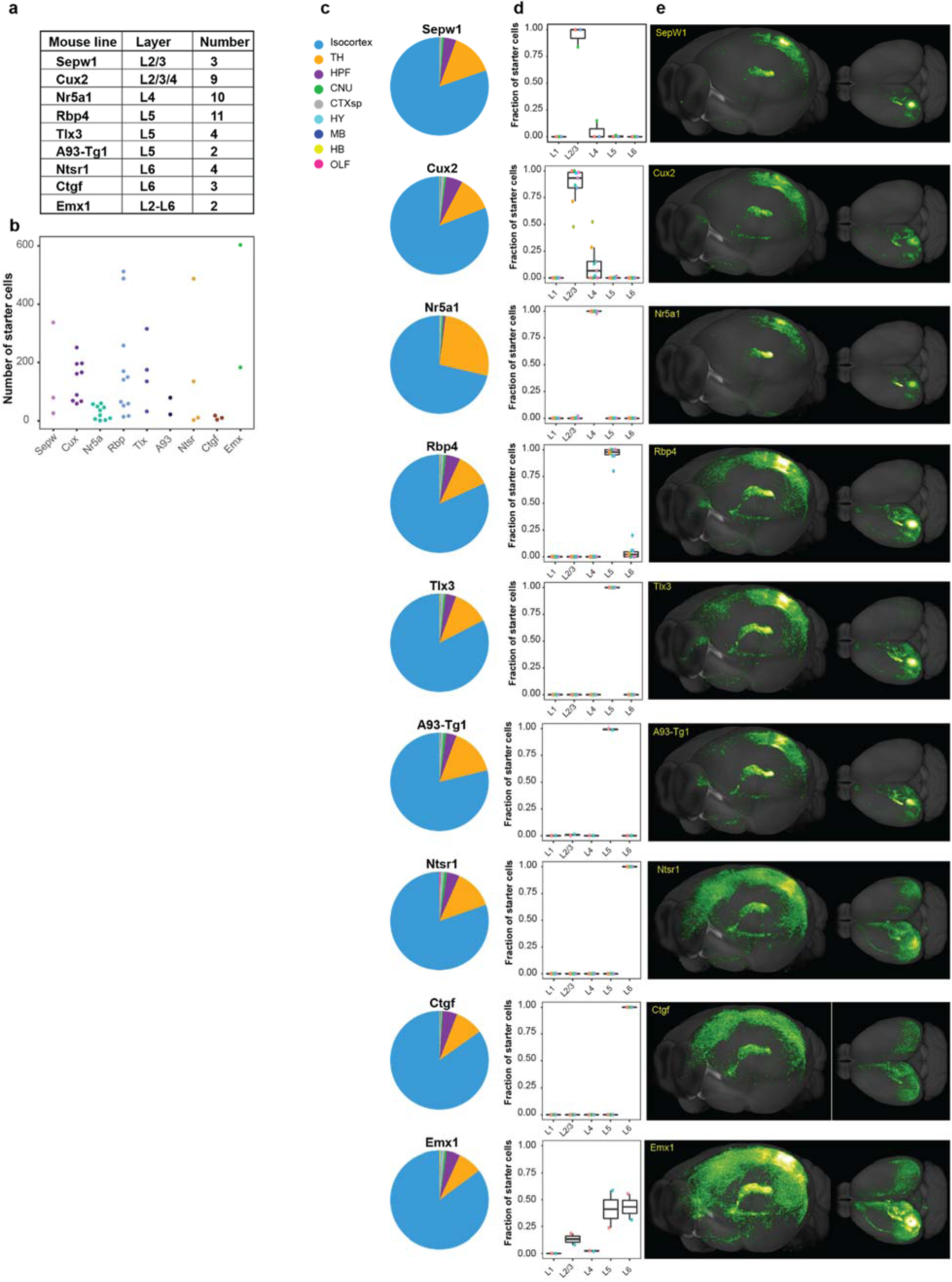
Whole-brain input patterns to excitatory neuron subclasses in different layers of the primary visual cortex. **(a-b)** Overview of the layer selectivity and number of experiments of each transgenic Cre line (a) and the numbers of starter cells grouped by Cre lines (b) for the 48 experiments in VISp. **(c)** Whole-brain input patterns of major brain regions to different layer-specific excitatory neuron subclasses labeled by the Cre lines. **(d)** Laminar distribution of starter cells for each Cre line. For each transgenic line, different experiments are indicated by different colors. **(e)** Representative 3D visualization of whole-brain inputs to neurons in different layers of VISp.

**Supplementary Figure 7.**
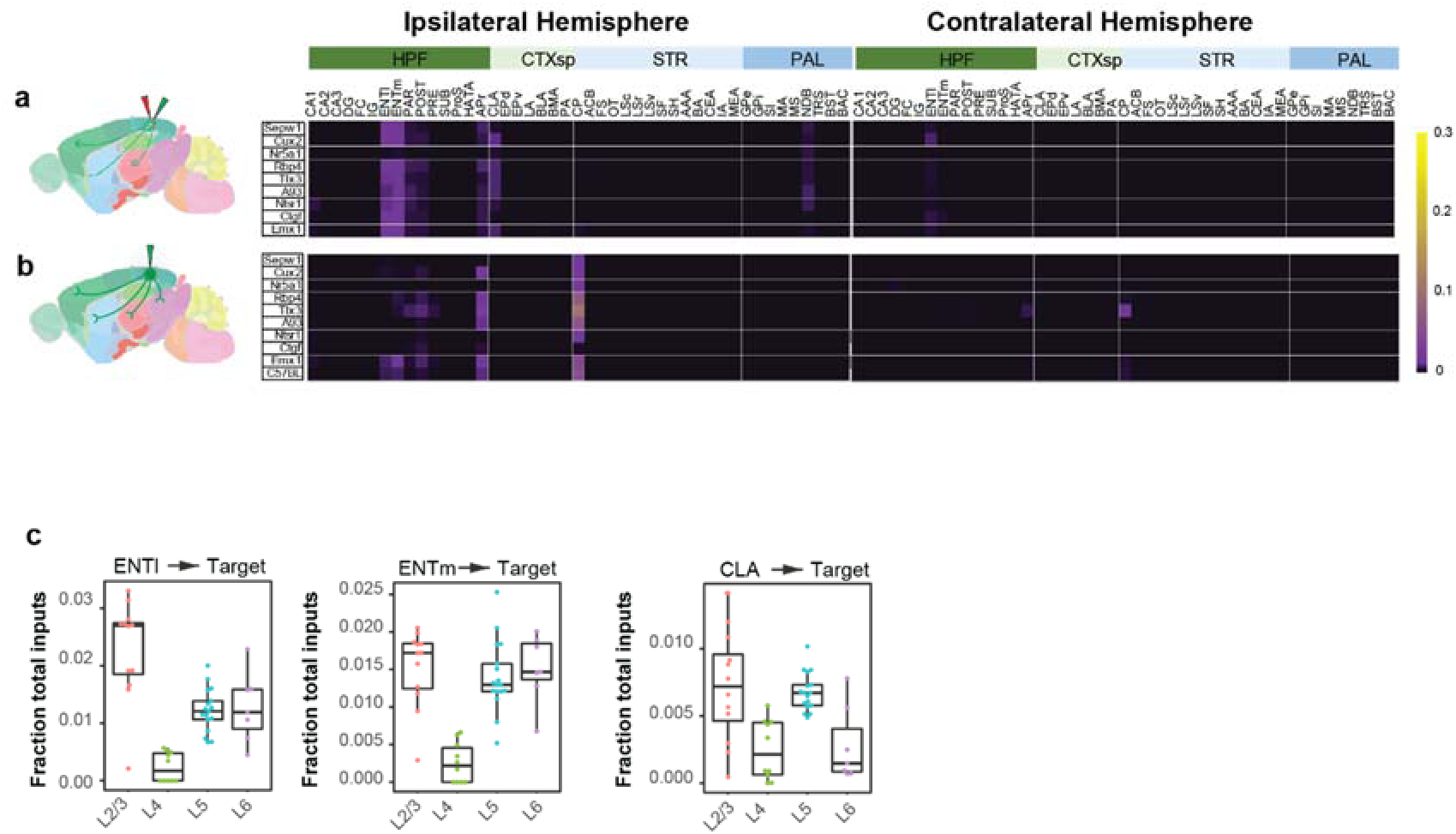
Comparison of subcortical inputs to excitatory neurons in different layers of the primary visual cortex. (**a)** Matrix showing presynaptic inputs from the ipsilateral and contralateral HPF, CTXsp, STR, and PAL to excitatory neurons in different layers of VISp. Each row of the matrix represents the mean normalized per structure input signals for experiments in each Cre line. Rows are organized based on layer-specific distribution of the starter cells. Brain areas are ordered by ontology order in the Allen CCF. **(b)** Matrix showing normalized projections from VISp to the brain regions shown in (a). Anterograde tracing experiments (**Supplementary Table 4**) from the Cre mouse lines used in (a) and C57BL/6J were included, and rows represent the mean normalized per structure projection signals for experiments in each mouse line. **(c)** Comparison of ENTl, ENTm, and CLA inputs to excitatory neurons in different layers of VISp.

**Supplementary Figure 8.**
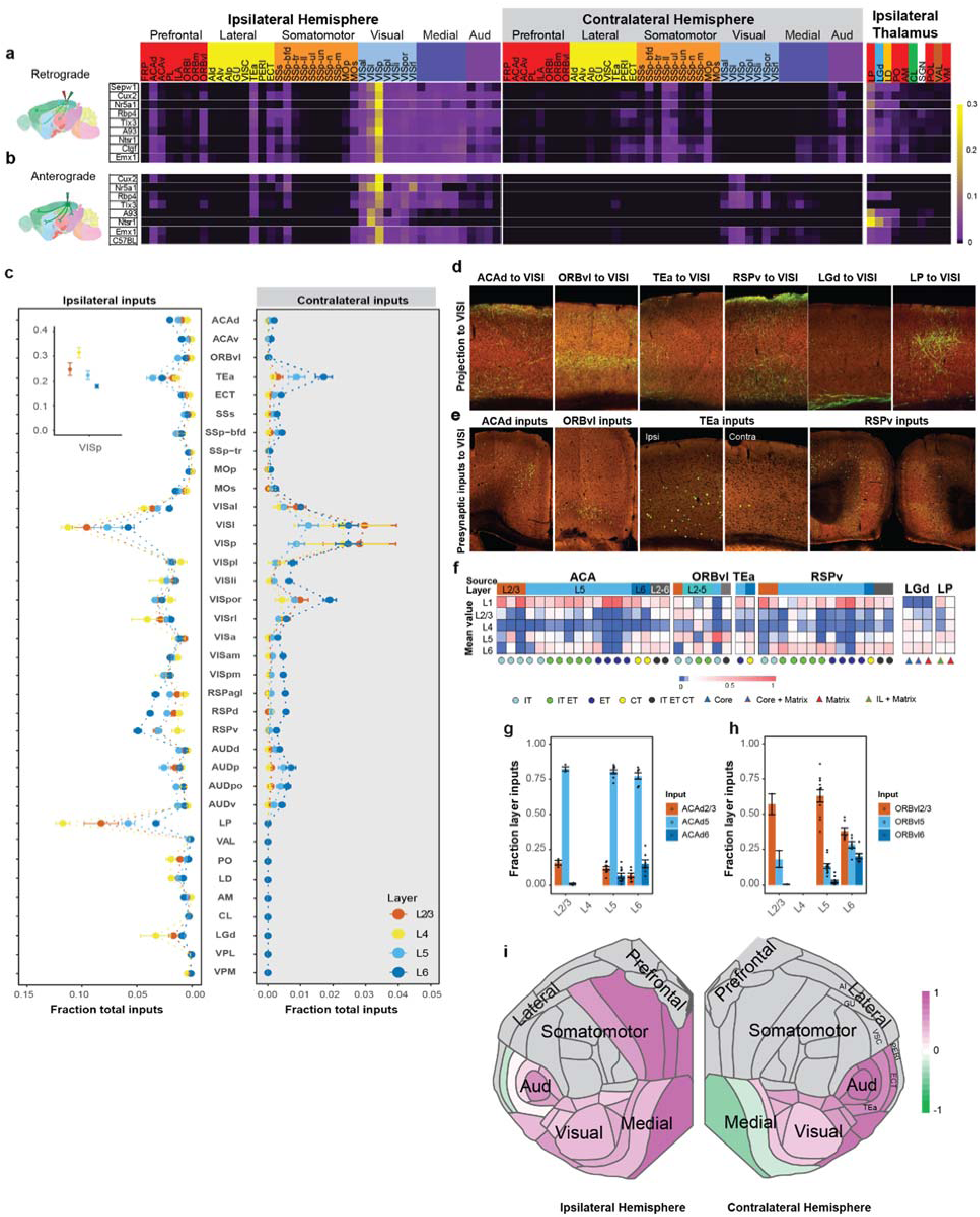
Comparison of the whole-brain input patterns to excitatory neuron subclasses in different layers of VISl. **(a)** Matrix showing normalized inputs from the ipsilateral and contralateral isocortex, and ipsilateral thalamus to excitatory neurons in different layers of VISl. Each row of the matrix represents the mean normalized per structure input signals for experiments in each Cre line. Rows are organized based on layer-specific distribution of the starter cells. The cortical areas are ordered first by module membership (color coded) then by ontology order in the Allen CCFv3. The ten thalamic nuclei are ordered based on the strength of inputs, and are color coded by the thalamocortical projection classes (blue: core, green: intralaminar, brown: matrix-focal, and red: matrix-multiareal). **(b)** Matrix showing normalized axon projections from VISl to the ipsilateral and contralateral isocortex, and ipsilateral thalamus shown in (a). Anterograde tracing experiments (**Supplementary Table 5**) from the Cre mouse lines used in (a) and C57BL/6J were included, and rows represent the mean normalized per structure projection signals for experiments in each mouse line. **(c)** Comparison of inputs from ipsilateral and contralateral cortical areas and thalamic nuclei to excitatory neurons in different layers of VISl. Data are shown as mean ± s.e.m. **(d-e)** Representative STPT images showing laminar termination patterns of axon projections in VISl from higher-order association cortical areas and thalamic nuclei (d) and laminar distribution patterns of presynaptic input cells in the cortical areas that project to VISl (e). **(f)** Normalized laminar termination patterns in VISl for projections from higher-order association cortical areas and thalamic nuclei. Each column represents the relative projection strengths by layer for a unique combination of Cre-defined cell classes and source areas. The average value was taken when n > 1. **(g-h)** Laminar distribution of long-range inputs from ACAd (g) and ORBvl (h) to excitatory neurons in different layers of VISl. The fraction layer input is calculated as the fraction of the total input signal in a given source area across layers. Data are shown as mean ± s.e.m. **(i)** Comparison of L5 and L6 input preference for source cortical areas in the ipsilateral (left) and contralateral (right) hemispheres sending presynaptic inputs to VISl. The preference score for a given cortical area is calculated as (L5 input - L6 input) / (L5 input + L6 input). Each source cortical area was colored according to its preference score.

**Supplementary Figure 9.**
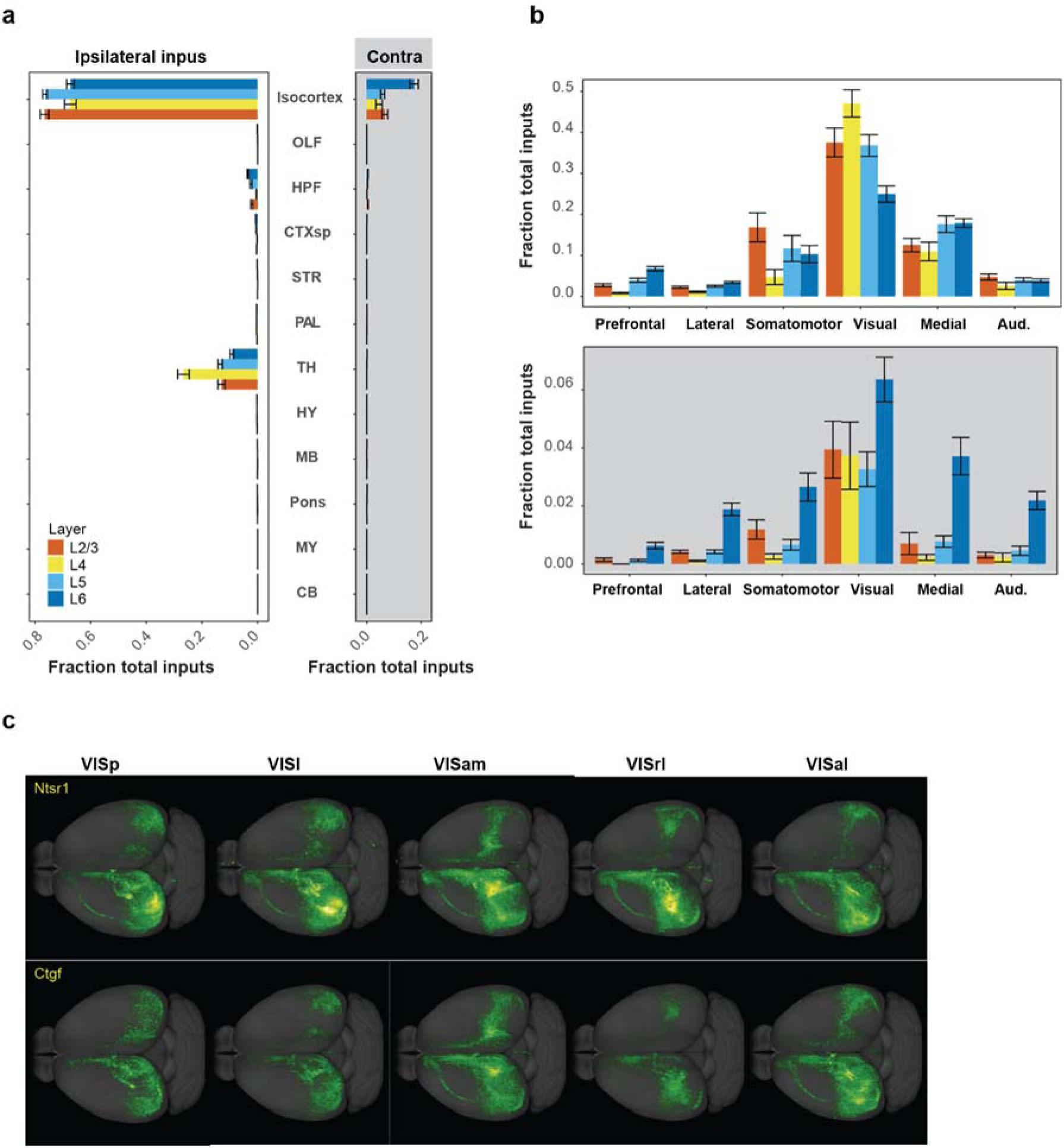
Comparison of contralateral cortical inputs to excitatory neuron subclasses in different layers of visual areas. **(a)** Comparison between ipsilateral (left) and contralateral (right) brain-wide inputs to excitatory neuron subclasses in L2/3, L4, L5 or L6 of visual areas. A total of 89 experiments with starter cells restricted in a single layer were identified, and the target areas included both VISp and HVAs. Data are shown as mean ± s.e.m. **(b)** Comparison between ipsilateral (top) and contralateral (bottom) isocortical inputs to excitatory neuron subclasses with starter cells restricted to either L2/3, L4, L5 or L6 of visual areas. Data are shown as mean ± s.e.m. **(c)** Representative top-down view of inputs to Ntsr1 and Ctgf-labeled L6 and L6b cell types in visual areas.

**Supplementary Figure 10.**
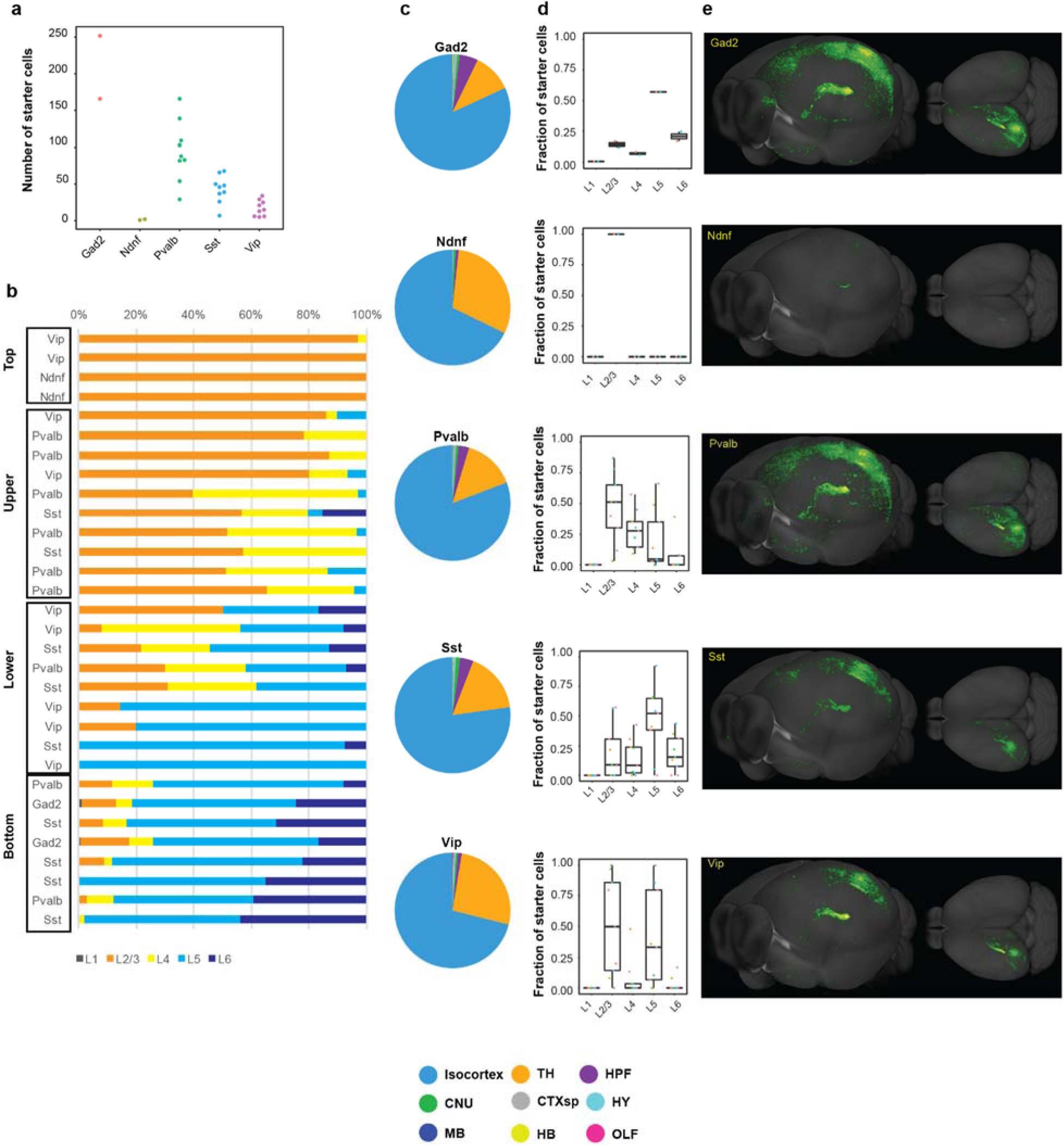
Analysis of the whole-brain input patterns to interneuron subclasses in the primary visual cortex. **(a)** Summary of the numbers of starter cells for each interneuron subclass. Each dot represents one individual experiment. **(b)** Fractions of starter cells located in L1, L2/3, L4, L5 and L6 for interneuron experiments in VISp. Experiments are divided into four groups based on distribution of the starter cells in different layers. **(c)** Overview of the whole-brain inputs to different interneuron subclasses. **(d)** Laminar distribution of starter cells for each interneuron subclass. For each transgenic line, different experiments are indicated by different colors. **(e)** Representative 3D visualization of whole-brain inputs to interneuron subclasses in VISp.

**Supplementary Figure 11.**
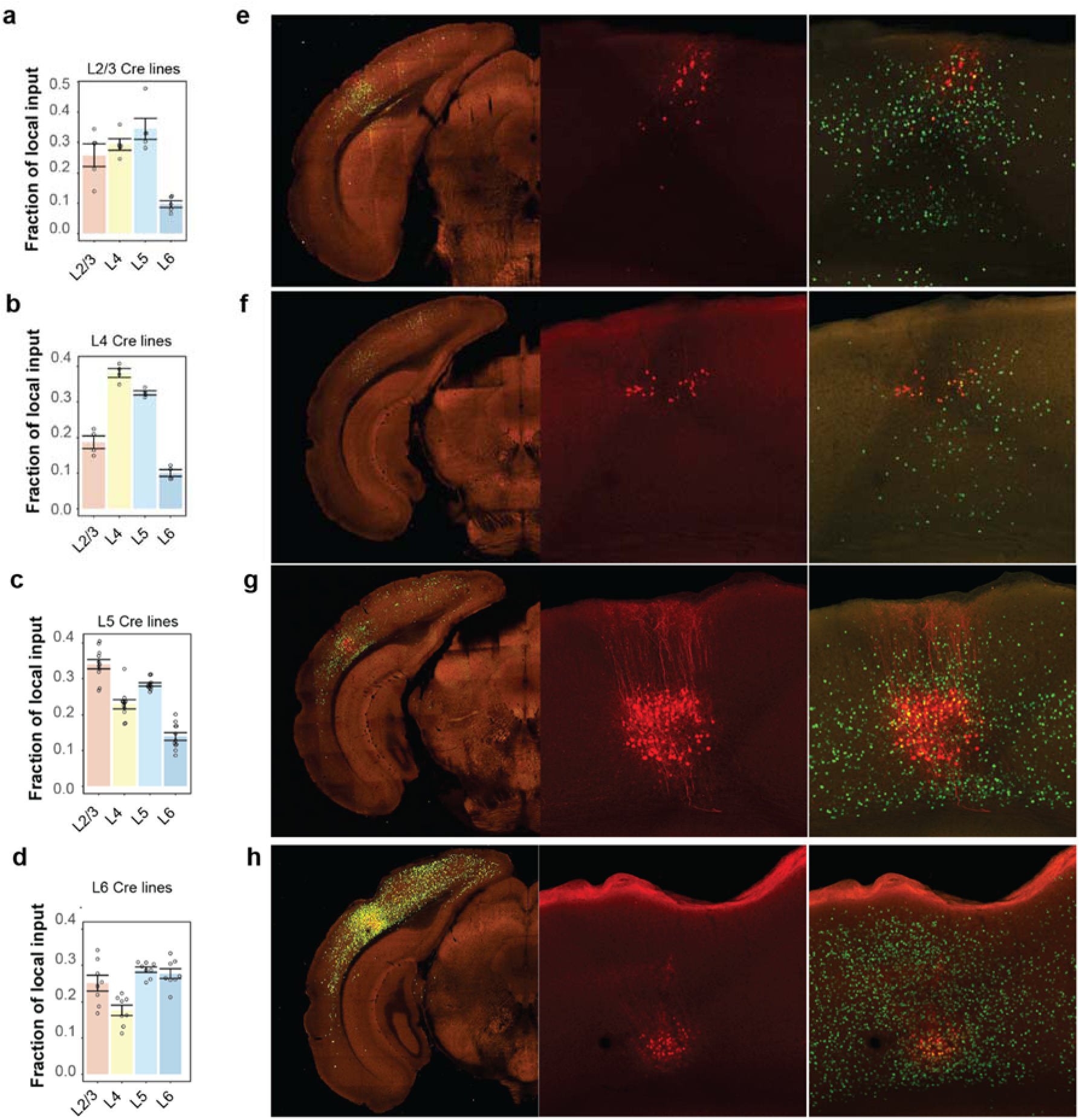
Comparison of local inputs to excitatory neurons in different layers of VISl. **(a-d)** Fraction layer input of ipsilateral VISl inputs to excitatory neurons in L2/3 (a), L4 (b), L5 (c) and L6 (d) of VISl. Data are shown as mean ± s.e.m. **(e-h)** Representative images showing layer-specific local inputs to excitatory neurons in L2/3 (e), L4 (f), L5 (g) and L6 (h) of VISl. Starter cells are identified by the coexpression of dTomato from the AAV helper virus and nucleus-localized EGFP from the rabies virus.

**Supplementary Figure 12.**
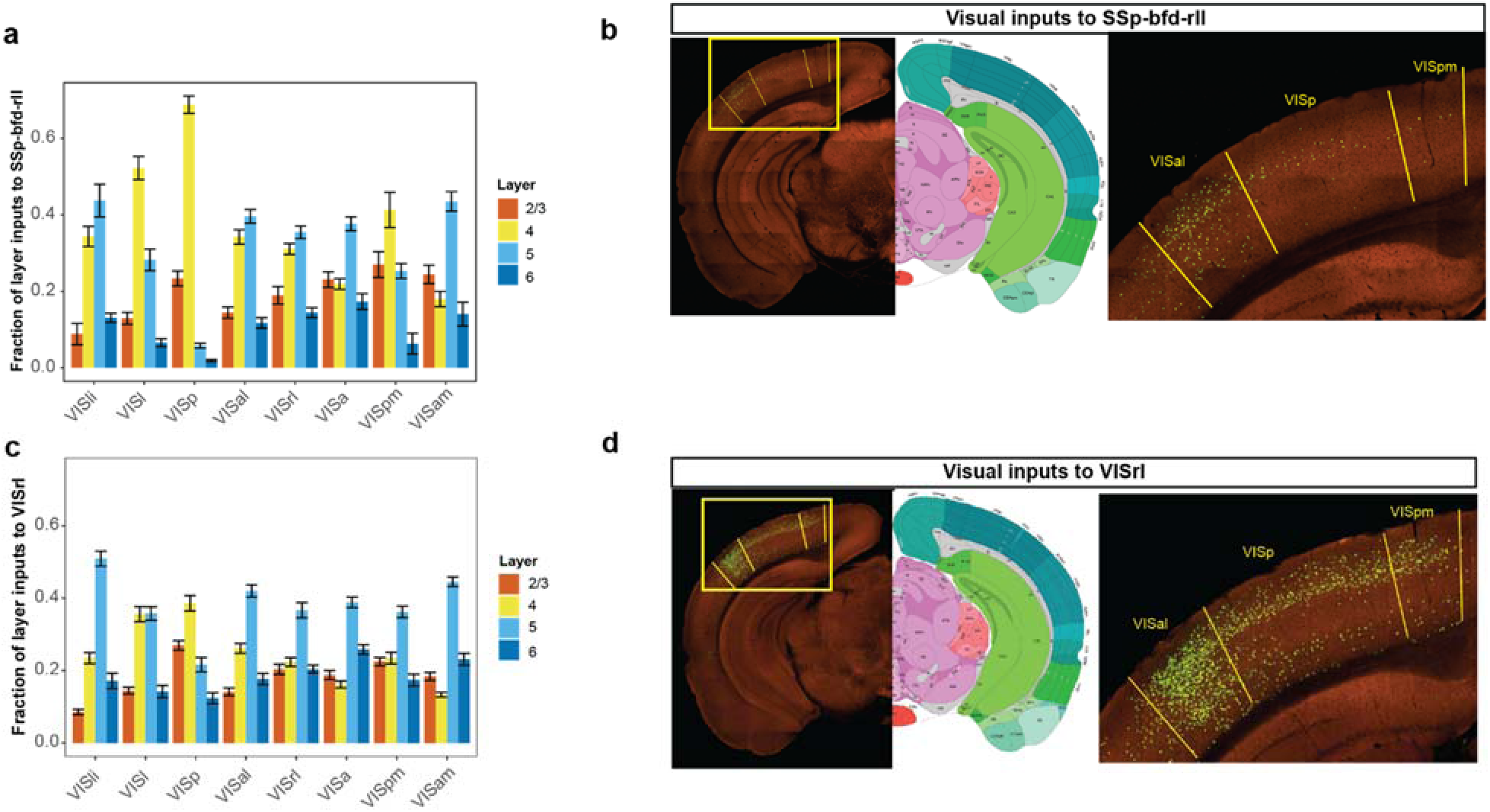
Comparison of laminar distribution of visual inputs to SSp-bfd-rll (a-b) and VISrl (c-d). **(a, c)** Laminar distribution of inputs from various visual areas to SSp-bfd-rll (a) and VISrl (c). Data are shown as mean ± s.e.m. **(b, d)** Representative images of inputs from VISp, VISpm and VISal to SSp-bfd-rll (b) and VISrl (d).

